# Systematic histone H4 replacement in *Arabidopsis thaliana* reveals a role for H4R17 in regulating flowering time

**DOI:** 10.1101/2022.01.17.476649

**Authors:** Emma Tung Corcoran, Chantal LeBlanc, Mia Arias Tsang, Anthony Sarkiss, Yuzhao Hu, Ullas V. Pedmale, Yannick Jacob

## Abstract

Despite the broad array of roles for epigenetic mechanisms on regulating diverse processes in eukaryotes, no experimental system for the direct assessment of histone function is currently available in plants. In this work, we present the development of a genetic strategy in *Arabidopsis thaliana* in which modified H4 transgenes can completely replace the expression of endogenous histone H4. Using this strategy, we established a collection of plants expressing different H4 point mutants targeting residues that may be post-translationally modified *in vivo*. To demonstrate the utility of this new H4 mutant collection, we screened it to uncover substitutions in H4 that alter flowering time. We identified different mutations in the tail (H4R17A) and the globular domain (H4R36A, H4R39K, H4R39A, and H4K44A) of H4 that strongly accelerate the floral transition. Furthermore, we found a conserved regulatory relationship between H4R17 and the ISWI chromatin remodeling complex in plants. Similar to other biological systems, H4R17 regulates nucleosome spacing via ISWI. Overall, this work provides a large set of H4 mutants to the plant epigenetics community that can be used to systematically assess histone H4 function in *A. thaliana* and a roadmap to replicate this strategy for studying other histone proteins in plants.

## Introduction

In eukaryotic cells, genomic DNA is organized into chromatin. The basic unit of chromatin is the nucleosome, which consists of 147 base pairs of DNA wrapped around a histone octamer made up of two copies of histone protein H2A, H2B, H3 and H4 (Luger et al., 1997). Histones play a significant role in regulating processes operating at the chromatin level, such as transcription and replication, and consequently, can have widespread effects on organismal growth, development and fitness (Kouzarides, 2007). One mechanism by which histones contribute to these processes is through the post-translational modifications (PTMs) of histone residues. Traditionally, the functional significance of histone PTMs has primarily been deduced through the analysis of phenotypes resulting from the mutation of histone-modifying enzymes or histone-reading proteins. However, while this method has been successful at identifying functions for many histone PTMs, there are several limitations to this approach. For example, this strategy presents difficulties if there are many redundant proteins writing or reading the same PTM, or if the writers or readers of a specific histone PTM have not been identified. Moreover, histone-modifying enzymes often target non-histone substrates in addition to histones, complicating the analysis of mutant phenotypes (Glozak et al., 2005).

To circumvent these obstacles, one effective strategy to study the functions of histone PTMs is to mutate the acceptor histone residue to an unmodifiable residue and then assess the resulting phenotype(s). One inherent advantage of this strategy is that it can be applied to assess the roles of modifiable and non-modifiable residues of histones. In addition, this histone replacement strategy can be used in biological backgrounds expressing wild-type histones, or in backgrounds where expression of endogenous histones is partially or completely eliminated. A major advantage of removing endogenous histones in this strategy is that it increases the likelihood of detecting phenotypes associated with expression of histone mutants, which may otherwise be masked if competing wild-type histones are also present. Systematic mutagenesis experiments with core histones were initially conducted in *Saccharomyces cerevisiae* to reveal many new insights into histone function (Dai et al., 2008; Fu et al., 2021; Govin et al., 2010; Nakanishi et al., 2008). These experiments utilized histone shuffle systems to either provide an episomal plasmid expressing histone mutants in a background with the endogenous histone genes deleted, or to directly mutate the endogenous histone copies using homologous recombination. The most recent system developed in *S. cerevisiae* utilized an efficient CRISPR/Cas9-based histone shuffle strategy that allows for the rapid development of multiplex histone mutations (Fu et al., 2021). In multicellular eukaryotes, *Drosophila melanogaster* is the first organism in which systematic histone mutagenesis was performed, and these systems used site-specific transgenesis to replace the endogenous histone coding region with that of a modified histone gene or histone array (Gunesdogan et al., 2010; Hodl and Basler, 2009, 2012; McKay et al., 2015). Additionally, high-throughput screens of histones H3 and H4 were recently conducted in this organism using a CRISPR/Cas9-mediated knock-in technology for histone gene replacement at the endogenous histone locus (Zhang et al., 2019).

In contrast to the aforementioned biological model systems, plant systems present additional challenges to implementing complete histone gene replacement. While all replication-dependent histone genes are clustered at a single genomic locus in *D. melanogaster* (Lifton et al., 1978), and *S. cerevisiae* contains only two copies of each core histone gene at the haploid cell stage (Fu et al., 2021), there are 47 genes encoding H2A, H2B, H3, and H4 in *Arabidopsis thaliana* found dispersed throughout the genome (Tenea et al., 2009). Because the histone genes are not clustered together in plants, the establishment of complex histone deletion mutants necessary for partially or completely replacing endogenous histone genes with modified histone genes is more challenging. Recently, histone replacement was performed in *A. thaliana* by using a combination of the traditional crossing of histone mutants and artificial microRNAs to generate backgrounds largely depleted of wild-type histone H3.1 (Jiang et al., 2017). However, this strategy is relatively time-consuming and may not completely eliminate endogenous histones. While the earliest strategies used to implement histone gene replacement in both *S. cerevisiae* and *D. melanogaster* were not applicable to plants due to their reliance on either the plasmid shuffle strategy and/or site-specific recombination systems, some aspects of the newest histone replacement strategies in other systems should facilitate the establishment of complete histone gene replacement in plants. For example, recent advancements in the deployment of multiplex CRISPR/Cas9-based technologies in plants make possible the creation of mutations in large gene families like histones.

While histones have been demonstrated to contribute to diverse processes in plants, a system enabling histone gene replacement would allow plant researchers to further elucidate the biological roles of histones in a more high-throughput manner. One of the most important developmental processes in plants is the transition from vegetative growth to reproductive development (Andres and Coupland, 2012; Song et al., 2015). Thus, the ways in which epigenetic mechanisms regulate the floral transition is a major area of research that could benefit from the application of histone replacement strategies. Diverse histone PTMs, including histone H3 lysine 4 (H3K4) methylation, H3K36 di- and tri-methylation, H3K9 methylation, H3K27 methylation, and H3 acetylation have been shown to regulate the expression of key flowering time regulatory genes such as *FLOWERING LOCUS C* (*FLC*) and *FLOWERING LOCUS T* (*FT*) (Bastow et al., 2004; Bu et al., 2014; Crevillen et al., 2019; Crevillen et al., 2014; Cui et al., 2016; Deng et al., 2007; He, 2009; He et al., 2004; Jiang et al., 2008; Kim et al., 2005; Ning et al., 2019; Pajoro et al., 2017; Pien et al., 2008; Xu et al., 2008; Yu et al., 2011; Zheng et al., 2019). However, compared to H3, the role of H4 in regulating the floral transition has been characterized to a much lesser extent.

Here, we present the establishment of a CRISPR-based histone mutagenesis platform in the plant model system *A. thaliana* that allows for complete histone replacement. As a proof-of-concept, we targeted histone H4, which is encoded by the largest number of endogenous genes (i.e. eight genes) among functionally-distinct histone proteins in plants (Okada et al., 2005; Tenea et al., 2009; Wierzbicki and Jerzmanowski, 2005), for systematic assessments of the roles of modifiable residues on this protein. After *in vivo* validation of our histone replacement strategy, we generated a large population of H4 point mutants to study the role(s) of 38 histone H4 residues. Using this H4 population, we identified a novel role for H4R17 in the regulation of flowering time. Furthermore, we demonstrated the functional relationship between H4R17 and an imitation switch (ISWI) chromatin-remodeling complex. Overall, this study demonstrates the utility of implementing histone replacement strategies in plants and provides a new resource that the plant community can use to probe for H4 functions in various aspects of plant growth and development.

## Results

### Generation of an *A. thaliana* mutant expressing a single histone H4 gene

In order to create a library of *A. thaliana* plants with replacement of endogenous histone H4 with H4 point mutants, we first generated a histone H4 depleted background using multiplex CRISPR/Cas9. The eight histone H4 genes in *A. thaliana* (Columbia [Col] ecotype) all code for the same histone H4 protein that is 98% identical to human histone H4 (100/102 identical a.a.; conservative substitutions at a.a. 60 and 77) (Supp. Fig. 1A). We designed three guide RNAs (gRNAs) that can target Cas9 to seven of the eight endogenous H4 genes (Supp. Fig. 1B). We then transformed Col plants via *Agrobacterium* using a multiplex Cas9/gRNA construct containing the three gRNAs against the H4 genes, selected first-generation transformants (T1), and exposed these T1 plants to repeated heat stress treatments at 37°C for 30h to increase the efficiency of targeted mutagenesis by Cas9 (LeBlanc et al., 2017). CRISPR/Cas9 activity was assessed at all seven H4 genes via PCR and sequencing in T1 plants, leading to the identification in the T2 generation of a plant with homozygous loss-of-function mutations in all seven targeted H4 genes (hereafter referred to as the H4 septuple mutant) (Supp. Fig. 1B). Morphological and molecular characterization of the H4 septuple mutant plants showed that they were slightly smaller than wild-type Col plants and displayed a serrated leaf phenotype (Fig. 1A). In addition, fertility was much lower in the H4 septuple mutant compared to Col plants (Fig. 1B). We found that transcription of the remaining endogenous H4 gene (*At3g53730*) was upregulated approximately 2-fold in the H4 septuple mutant relative to Col, likely to compensate for H4 depletion due to the loss of function of the other seven H4 genes (Fig. 1C). The H4 septuple mutant exhibited mis-regulation of markers of genomic and epigenomic instability, including up-regulation of the DNA damage response gene *BRCA1* and transcriptional de-repression of the heterochromatic DNA repeat *TSI* (Fig. 1D-E). These results demonstrate that multiplex CRISPR/Cas9 can be used to rapidly create an *A. thaliana* mutant background containing a minimal amount of functional genes coding for a specific histone.

**Figure 1:**
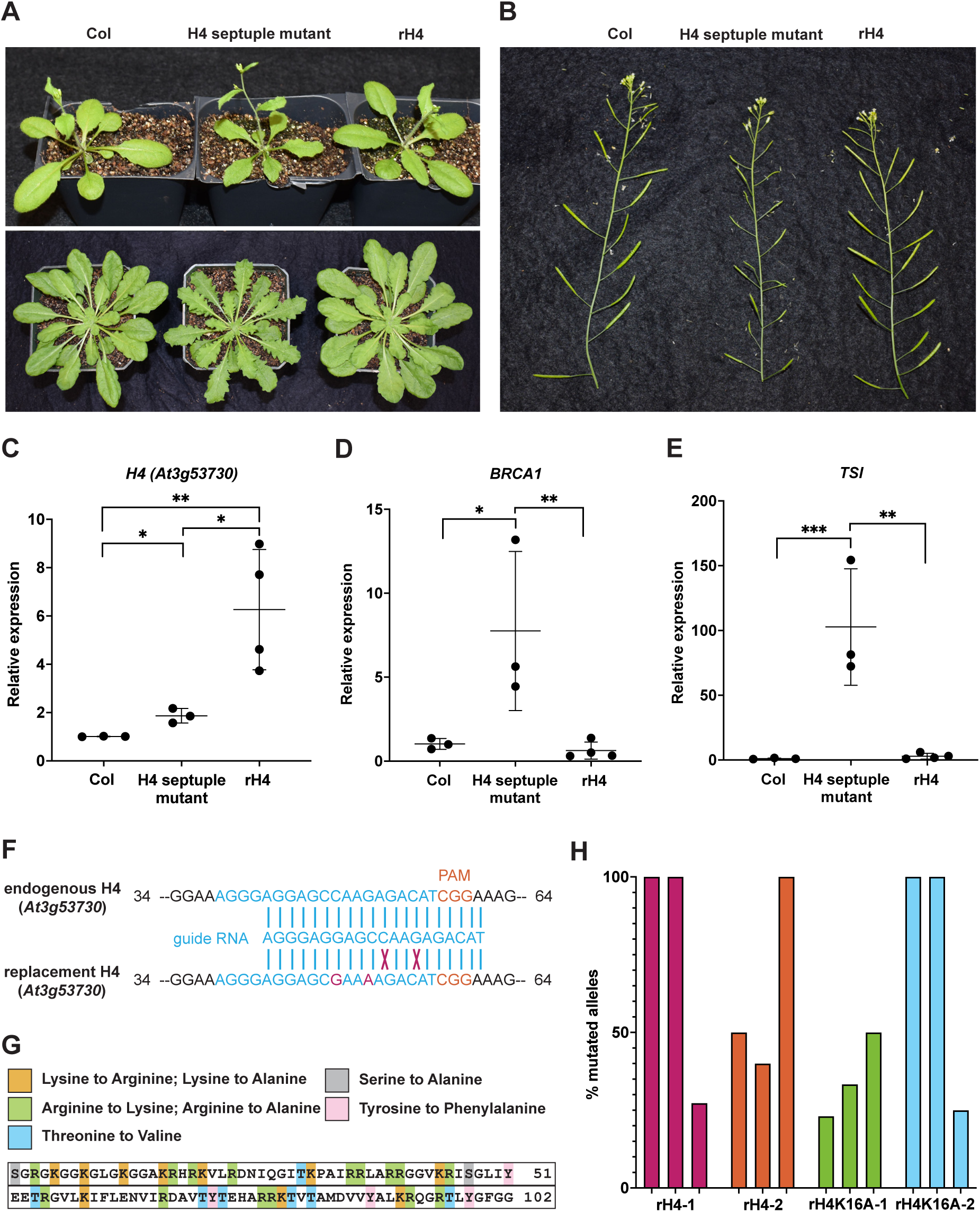
A CRISPR-based genetic system for expression of H4 point mutants in *A. thaliana*. (A) Morphological phenotypes of Col, H4 septuple mutants, and rH4 plants grown in long-day conditions at 3.5 weeks (top) and short-day conditions at 7 weeks (bottom). (B) Siliques of Col, H4 septuple mutants, and rH4 plants grown in long-day conditions for 4 weeks. (C-E) RT-qPCR of (C) *H4* (*At3g53730*), (D) *BRCA1* and (E) *TSI* in Col, H4 septuple mutants, and four independent rH4 T1 lines. Three biological replicates were included for Col and H4 septuple mutant plants. Horizontal bars indicate the mean. SD denoted with error bars. *P*-value from unpaired Student’s *t*-test denoted with asterisks (*p<0.05, **p<0.005, ***p<0.0005). (F) Design of the gRNA targeting the remaining endogenous H4 gene (*At3g53730*) in the H4 septuple mutant. Mismatches of the replacement H4 gene with the gRNA shown in red. (G) Schematic of point mutations in the H4 replacement plasmid library. (H) Percentage of mutated alleles in six rH4 plants and six rH4K16A plants. Each plant assessed was from the T2 generation; three plants from the same T1 parent were used in this experiment (i.e. two independent T1 lines per genotype).

### Establishment of a histone H4 replacement system in *A. thaliana*

To set up complete histone H4 replacement in plants, we designed an H4 replacement plasmid that contains 1) a gRNA targeting the last remaining endogenous H4 gene (*At3g53730*) and 2) a Cas9-resistant H4 gene allowing for expression of *At3g53730* under its native promoter (i.e. H4 replacement gene). Our strategy was to transform the H4 septuple mutant, which expresses Cas9, with the H4 replacement plasmid and select T1 plants that contain mutations at the endogenous *At3g53730* gene. To prevent Cas9 from targeting the replacement H4 gene, we introduced two silent mutations in this gene that prevent recognition from the gRNA targeting the endogenous *At3g53730* gene (Fig. 1F). After transformation of the H4 septuple mutant with the H4 replacement plasmid, we recovered many T1 transformants expressing the replacement H4 gene (hereafter referred to as rH4 plants), and in contrast to the H4 septuple mutant, all rH4 plants were normal in size, did not exhibit serrated leaves and showed normal fertility (Fig. 1A-B). Moreover, the RNA expression levels of *BRCA1* and *TSI* in rH4 plants were comparable to levels observed in Col (Fig. 1D-E). The expression of *At3g53730* in first-generation rH4 plants was found to be upregulated approximately 4- to 9-fold relative to Col (Fig. 1C). These results indicate that high expression levels of the replacement H4 gene in rH4 plants are responsible for suppressing the morphological phenotypes of the H4 septuple mutant.

We then used site-directed mutagenesis to create a large library of H4 replacement plasmids carrying different point mutations in the H4 replacement gene. We generated mutations covering every amino acid (i.e. lysine, arginine, threonine, serine, and tyrosine) in H4 that could theoretically be post-translationally modified *in vivo*. Each modifiable amino acid was mutated to a residue that cannot be post-translationally modified (i.e. alanine, valine or phenylalanine). We also substituted lysine and arginine residues with residues having similar biochemical properties (i.e. arginine and lysine, respectively). In total, we modified 38 amino acid residues of H4 to generate 63 different H4 replacement genes containing a specific point mutation (Fig. 1G). We subcloned these H4 mutant genes into the H4 replacement plasmid and individually transformed them into the H4 septuple mutant. We selected two independent transgenic lines for each H4 mutant, except for plants expressing the replacement genes H4 arginine 40 to alanine (rH4R40A), rH4R45A, rH4K59A, rH4R78A, rH4K79R and rH4R92K due to lethality induced by these mutations. All T1 plants were exposed to repeated heat stress treatments to maximize the efficiency of targeted mutagenesis of the remaining endogenous H4 gene by Cas9. To estimate the frequency of mutations at the remaining endogenous H4 gene in the plants expressing the replacement H4 gene, we genotyped three plants each from two independent rH4 lines and two independent rH4K16A lines at the T2 generation stage. We amplified the remaining endogenous H4 gene (*At3g53730*) from these T2 plants, cloned the resulting PCR products and sequenced at least ten individual clones corresponding to each plant, and calculated the percentage of mutated alleles. Approximately half of the plants were characterized by a complete elimination of the wild-type *At3g53730* allele, while the other plants varied from 50 to 75% wild-type alleles remaining (Fig. 1H). Taking into account that expression of the replacement H4 gene is either equivalent or much higher compared to the remaining endogenous H4 gene (Fig. 1C), these results suggest that the chromatin of most T2 plants in our H4 replacement collection contains large amounts of H4 point mutants. Overall, these results show that our CRISPR strategy was successful in creating a large collection of *A. thaliana* plants expressing different H4 point mutants replacing wild-type H4 proteins.

### Differential regulation of flowering time in plants expressing histone H4 mutants

To demonstrate the utility of the H4 replacement collection in identifying pathways regulated by H4 in *A. thaliana*, we initiated a screen of the plants expressing H4 mutants for defects in flowering time. The transition between vegetative and reproductive development in *A. thaliana* has been shown to be sensitive to various chromatin disruptions, but most of the findings in this field have focused on the roles of post-translational modifications on histone H3 (Berry and Dean, 2015; He, 2009; He and Amasino, 2005; Srikanth and Schmid, 2011; Yaish et al., 2011).

We grew our collection of H4 mutants using T2 plants from two independent lines for each H4 mutant, and measured flowering time (days to flowering and leaf number) in both short-day conditions (8 hours light, 16 hours dark) and long-day conditions (16 hours light, 8 hours dark). We identified many morphological and developmental phenotypes at the vegetative stage of growth in T2 plants expressing the different H4 mutants (Fig. 2A, Supp. Fig. 2A), which demonstrates that our H4 replacement strategy can be used to reveal various developmental phenotypes associated with mutations on histone H4. In regard to flowering time, we did not observe plants expressing H4 point mutants that were associated with a consistent and significant late flowering-time phenotype compared to rH4 plants (i.e. replacement with WT H4 gene) for both transgenic lines corresponding to the same mutation in either long-day or short-day conditions. In contrast, 16 rH4 mutants were identified as displaying an early flowering phenotype using the same criteria (Fig. 2B-C, Supp. Fig. 2B-C, Supp. Fig. 3). Many rH4 mutant lines exhibited early flowering in both long-day and short-day conditions, with the rH4R17A, rH4R36A, rH4R39K, rH4R39A, and rH4K44A mutants exhibiting the most consistent and drastic decrease in flowering time. In order to reduce the dimensionality of the data, we performed principal component analysis of the four flowering time variables measured in our analyses: mean day number in long-day, mean leaf number in long-day, mean day number in short-day, and mean leaf number in short-day (Fig. 2D). We performed *k*-means clustering on PC1 and PC2, which together explained 98% of the variance (Supp Fig. 2D), to identify three clusters in the data. Cluster a, corresponding to a flowering response most similar to wild-type plants, contained Col, the H4 septuple mutant, rH4, rH4K16A, rH4K20R, and rH4K20A. Cluster b, corresponding to a moderately early flowering time phenotype, contained rH4R17K, rH4R35K, rH4R35A, rH4R36K, rH4R40K, and rH4K44R. Cluster c, corresponding to a drastically early flowering time phenotype, contained rH4R17A, rH4R36A, rH4R39K, rH4R39A, and rH4K44A. The two rH4K16R lines were split between Cluster a and Cluster b, and the two rH4T80V lines were split between Cluster b and Cluster c. While the rH4K16R, rH4K16A, rH4K20R, and rH4K20A mutants appeared slightly early flowering relative to rH4 plants (Fig. 2B-C, Supp. Fig. 2B-C), all of these mutant lines except for a single rH4K16R line clustered within the wild-type cluster (Cluster a) through these analyses. We performed RT-qPCR analyses on key genes (*FLOWERING LOCUS T* (*FT*) and *SUPPRESSOR OF OVEREXPRESSION OF CO 1* (*SOC1*)) regulating flowering time and observed upregulation of these genes consistent with the early flowering behavior of rH4 mutants from different clusters (Fig. 2E-F). Taken together, our histone H4 replacement system enables the assessment of expressing histone H4 mutants on flowering time regulation, thus demonstrating the usefulness of the system for probing histone H4 function in plants.

**Figure 2:**
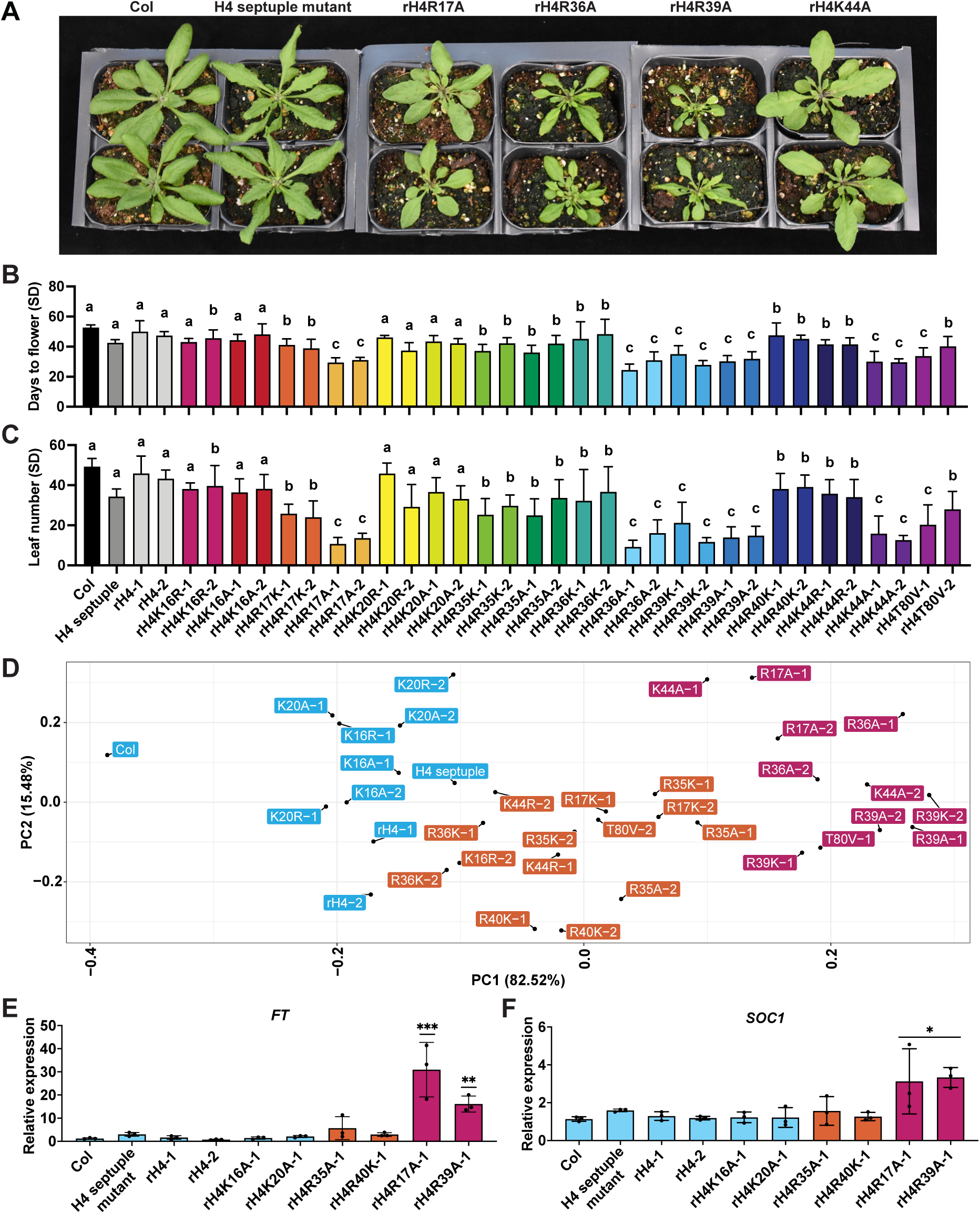
Mutations in specific residues of histone H4 generate early flowering phenotypes in *A. thaliana*. (A) Rosette phenotype of Col, H4 septuple mutant, rH4R17A, rH4R36A, rH4R39A, and rH4K44A mutants grown in long-day conditions for 3 weeks. For the rH4 plants, individual T2 plants (top and bottom) from independent T1 parents are shown. (B-C) Mean (B) days to flower and (C) rosette leaf number at flowering in short-day (SD) conditions for Col, the H4 septuple mutant, and various H4 replacement backgrounds (two independent transgenic lines each). Standard deviation shown with error bars (n≥7). Letters (a,b,c) indicate cluster identified by *k*-means clustering. (D) Principal component plot for flowering time data along the first two principal components, PC1 and PC2. Variance explained by each principal component shown on respective axis. Three clusters produced by *k*-means clustering represented in blue (Cluster a), orange (Cluster b), and pink (Cluster c) colors. (E-F) RT-qPCR of (E) *FLC* and (F) *SOC1* in Col, H4 septuple mutant, rH4-1, rH4-2, rH4K16A-1, rH4K20A-1, rH4R35A-1, rH4R40K-1, rH4R17A-1, and rH4R39A-1 plants. Standard deviation denoted with error bars. Statistical analyses were performed using one-way ANOVA with Tukey’s HSD post hoc test. *P*-value from Tukey’s HSD test (genotype vs. Col) denoted with asterisks (*p<0.05, **p<0.005, ***p<0.0005). Bar colors represent cluster assignment from (D).

**Figure 3:**
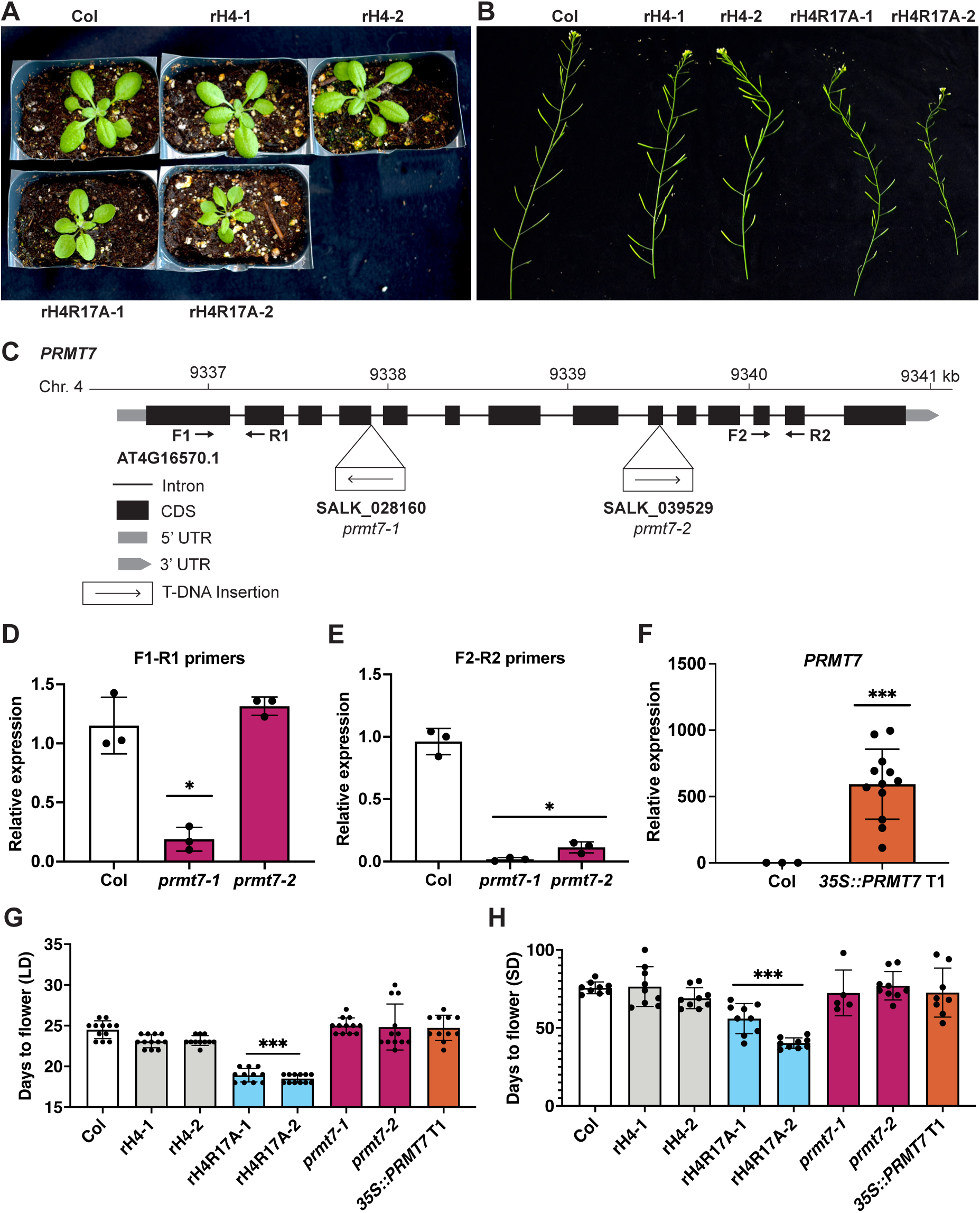
*PRMT7* does not regulate the floral transition in *A. thaliana*. (A) Rosette phenotype of Col, rH4-1, rH4-2, rH4R17A-1 and rH4R17A-2 plants grown in long-day conditions at 3 weeks. (B) Silique phenotype of Col, rH4-1, rH4-2, rH4R17A-1 and rH4R17A-2 plants grown in long-day conditions at 4 weeks. (C) Gene structure of *PRMT7*. The location of the T-DNA insertions and the primers (F1-R1 and F2-R2) used for gene expression analyses are shown. (D-F) RT-qPCR showing *PRMT7* expression in (D-E) Col, *prmt7-1*, and *prmt7-2* plants and (F) Col and *35S::PRMT7* T1 plants. The average of three biological replicates and standard deviation are shown for Col and *prmt7* mutants. For the *35S::PRMT7* plants, individual data points represent independent T1 plants. *P*-value from unpaired Student’s *t*-test (sample vs. Col) denoted with asterisks (*p<0.05, **p<0.005, ***p<0.0005). (E-F) Mean days to flower in (E) long-day (LD) and (F) short-day (SD) conditions for Col, rH4-1, rH4-2, rH4R17A-1, rH4R17A-2, *prmt7-1, prmt7-2,* and *35S::PRMT7* T1 plants. Standard deviation denoted with error bars. Statistical analyses were performed using one-way ANOVA with Tukey’s HSD post hoc test. *P*-value from Tukey’s HSD test (genotype vs. Col) denoted with asterisks (*p<0.05, **p<0.005, ***p<0.0005). n≥11 for long-day, n≥5 for short-day.

### *In vivo* modulation of *PRMT7* activity does not replicate the early flowering phenotype of rH4R17A plants

Among the five H4 mutations (H4R17A, H4R36A, H4R39K, H4R39A, and H4K44A) identified in our screen to cause the strongest effect on flowering time, only one of them (H4R17A) is present in the N-terminal tail (a.a. 1-20) of H4 where most of the histone post-translational modifications (PTMs) are made. Mutations in the unstructured N-terminal tail of H4 are less likely to affect flowering time by disrupting histone H4 folding and/or nucleosome structure than mutations in the histone-fold domain. Therefore, we focused our subsequent analyses on trying to elucidate the mechanism by which H4R17A affects the timing of the transition to reproductive development. For this work, we used H4 replacement plants for which there was a complete replacement of the endogenous histone H4 with H4R17A (Supp. Fig. 4A). In addition to a significantly early floral transition, we found that the H4R17A mutation also causes other developmental phenotypes, including smaller, upwardly curled leaves and reduced fertility compared to wild-type plants (Fig. 3A-B).

**Figure 4:**
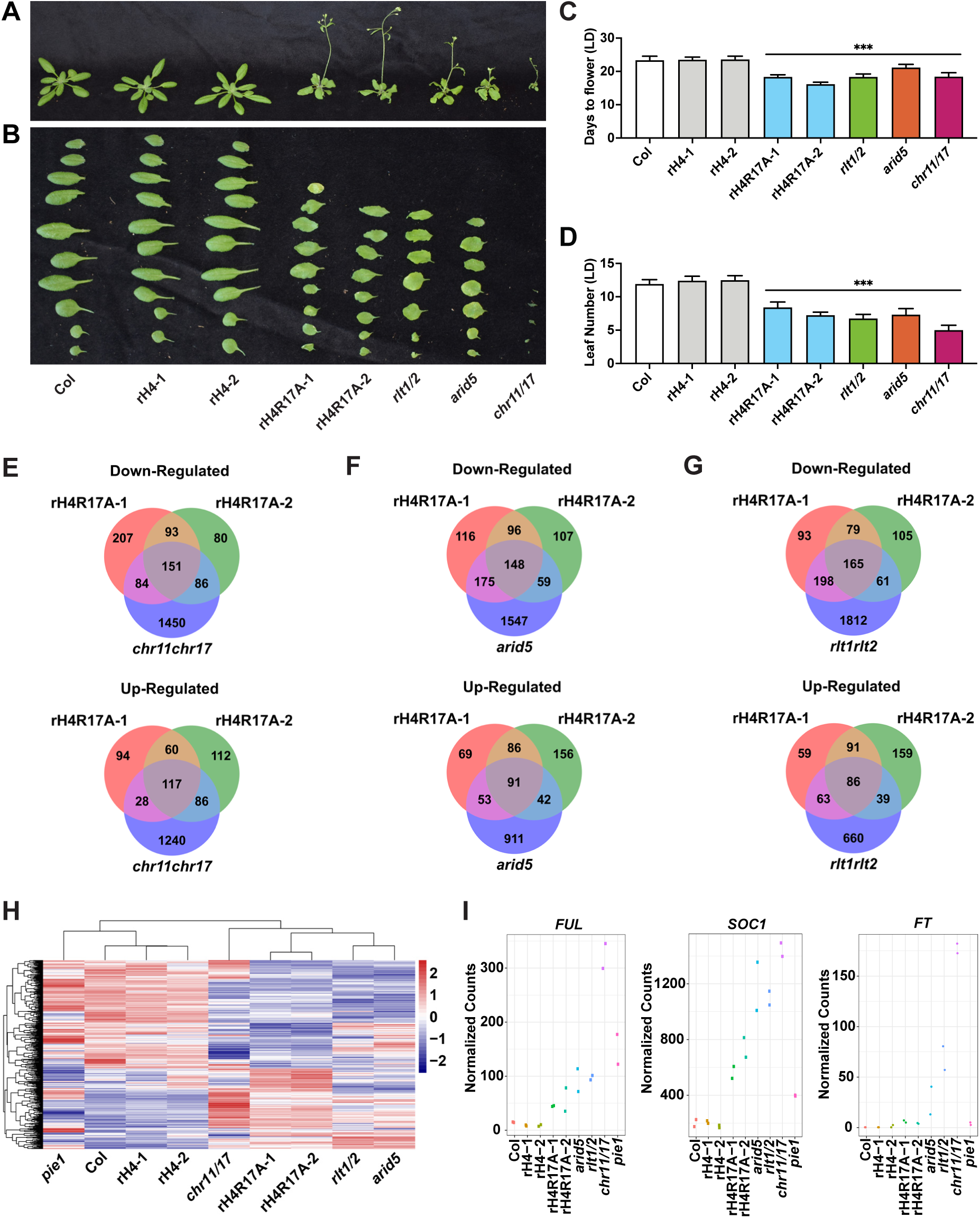
H4R17 and ISWI are functionally related in the regulation of gene expression and plant development. (A) Morphological phenotypes of Col, rH4-1, rH4-2, rH4R17A-1, rH4R17A-2, *rlt1/2, arid5*, and *chr11/17* plants grown in long-day conditions at 21 days. (B) Rosette leaf phenotype of Col, rH4-1, rH4-2, rH4R17A-1, rH4R17A-2, *rlt1/2, arid5*, and *chr11/17* plants. Rosette leaves were cut from plants shortly after bolting in long-day conditions. (C-D) Mean (C) days to flower and (D) rosette leaf number at flowering in long-day (LD) conditions for Col, rH4-1, rH4-2, rH4R17A-1, rH4R17A-2, *rlt1/2, arid5*, and *chr11/17* plants. Standard deviation shown with error bars. Statistical analyses were performed using one-way ANOVA with Tukey’s HSD post hoc test. *P*-value from Tukey’s HSD test (genotype vs. Col) denoted with asterisks (*p<0.05, **p<0.005, ***p<0.0005). n=12. (E-G) Venn diagrams showing DEGs (relative to Col) identified by RNA-seq in (E) rH4R17A and *chr11/17*, (F) rH4R17A and *arid5*, and (G) rH4R17A and *rlt1/2* plants. (H) Heatmap of relative expression patterns of shared DEGs identified in the rH4R17A-1 and rH4R17A-2 mutants. Legend represents scaled Z-score on normalized read counts. Clustering of rows and columns calculated using Euclidean distance. (I) Normalized read counts at *FUL*, *SOC1*, and *FT* in Col, rH4-1, rH4-2, rH4R17A-1, rH4R17A-2, *arid5, rlt1/2, chr11/17*, and *pie1* plants.

One hypothesis regarding the mechanism by which the H4R17A mutation causes early flowering is that it prevents deposition of a post-translational modification on H4R17. PROTEIN ARGININE METHYLTRANSFERASE 7 (PRMT7) is the only known histone-modifying enzyme of H4R17 in eukaryotes, as it has been shown to mono-methylate H4R17 in mammals (Feng et al., 2014; Feng et al., 2013; Jain and Clarke, 2019). The *A. thaliana* genome contains a single orthologous gene for *PRMT7* (*At4g16570*), which has never been functionally characterized. To assess a potential role for PRMT7 in regulating flowering time via methylation of H4R17, we measured flowering time in *prmt7* mutants (SALK_028160 and SALK_039529) and in plants overexpressing the *PRMT7* gene (i.e. *35S::PRMT7*). We confirmed by RT-qPCR that both T-DNA alleles used in these experiments prevent the expression of a full-length *PRMT7* transcript and that *PRMT7* was overexpressed in the *35S::PRMT7* plants that we generated (Fig. 3C-F). Our analyses of flowering time caused by modulation of the *PRMT7* gene in plants showed that neither *prmt7* mutants nor *PRMT7* overexpressing plants displayed altered flowering time in either long-day conditions or short-day conditions (Fig. 3G-H, Supp. Fig. 4B-C). In addition, none of the other vegetative or reproductive phenotypes observed in rH4R17A plants were found in plants lacking or overexpressing *PRMT7*. These results strongly suggest that replacement of H4 with H4R17A does not affect development in *A. thaliana* by interfering with PRMT7 activity on histone H4.

### Functional relationship between H4R17 and ISWI in the regulation of flowering time

In addition to affecting the deposition of PTMs, mutation of histone residues can prevent binding of proteins to chromatin (Hyland et al., 2005; Norris et al., 2008). Therefore, we next investigated the possibility that replacement of histone H4 with H4R17A affects plant development by negatively impacting the function of plant ISWI chromatin-remodeling complexes. In yeast and animals, R17 of H4 has been shown to directly interact with ISWI to regulate nucleosome remodeling activity *in vitro* and *in vivo* (Clapier et al., 2001; Clapier et al., 2002; Dann et al., 2017; Fazzio et al., 2005; Hamiche et al., 2001; Ludwigsen et al., 2017; Mueller-Planitz et al., 2013; Racki et al., 2014; Yan et al., 2016). Mutations in genes coding for different *A. thaliana* ISWI subunits (CHR11, CHR17, RLT1, RLT2 and ARID5) result in plants showing similar phenotypes as rH4R17A mutants, including early flowering, upwardly curled leaves, reduced fertility, and a small size relative to wild-type plants (Fig. 4A-D, Supp. Fig. 5) (Li et al., 2012). Defects in the timing of the floral transition and other developmental aspects are more similar between the rH4R17A mutants and the ISWI accessory subunit single mutant *arid5* and double mutant *rlt1 rlt2* (*rlt1/2*; *RLT1* and *RLT2* were shown to act redundantly (Li et al., 2012)) compared to mutations in the ISWI catalytic subunits *CHR11* and *CHR17* (*CHR11/17*; also shown to act redundantly (Li et al., 2012)), which cause more severe developmental phenotypes (Fig. 4A-D, Supp. Fig. 5) (Li et al., 2012). The increased severity of the phenotypes displayed by the *chr11/17* double mutant may be caused by the joint disruption of the ISWI and SWR1 chromatin-remodeling complexes, which both contain CHR11/17 (Luo et al., 2020). In contrast, ARID5 and RLT1/2 are present in ISWI, but not in SWR1. In addition, RLT1 and RLT2 are only two of 12 DDT-domain proteins in *A. thaliana*, and different ISWI complexes were found to associate with different DDT-domain proteins *in vivo* (Dong et al., 2013; Tan et al., 2020).

**Figure 5:**
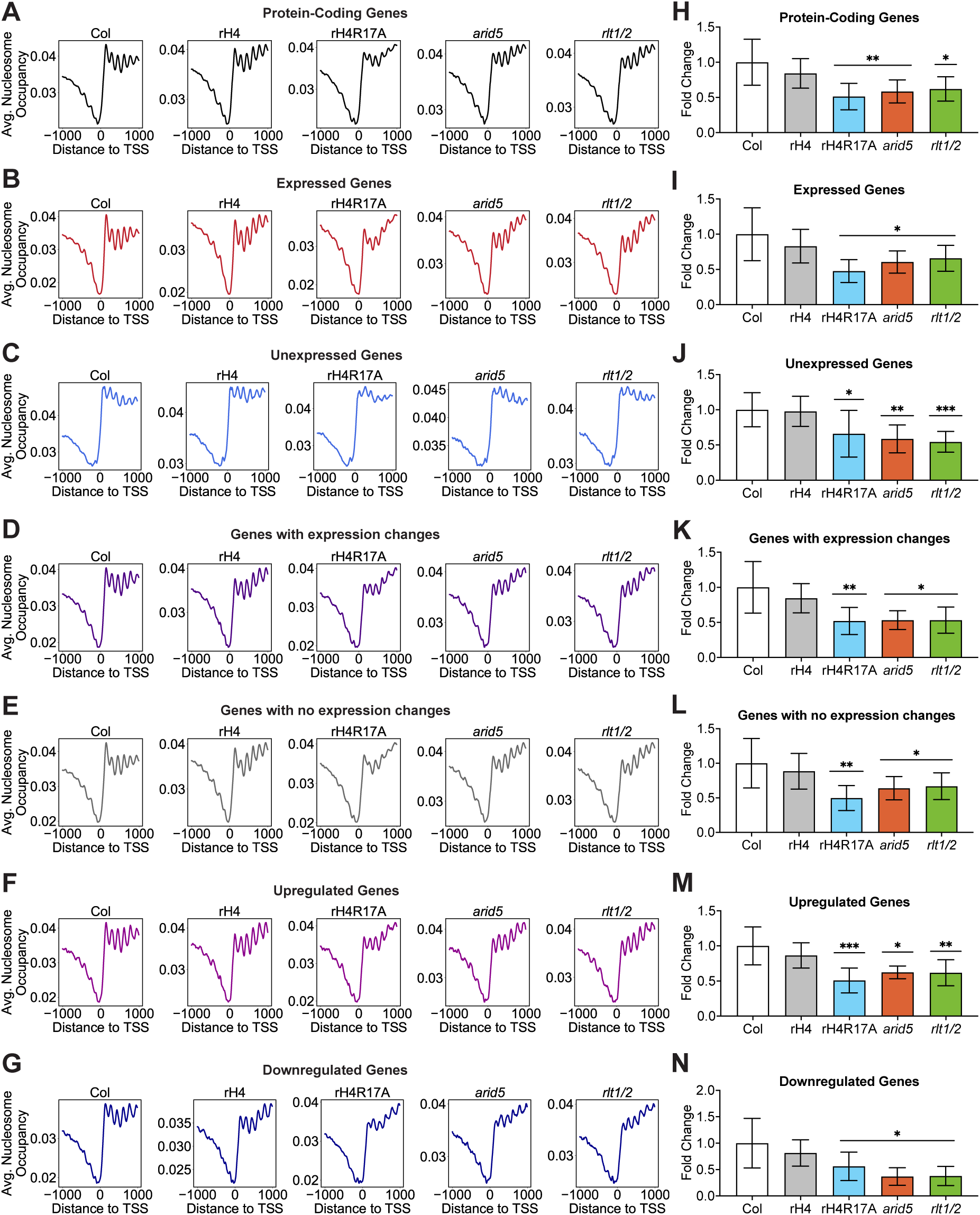
Determination of the function of H4R17 on regulating nucleosome positioning. (A-G) Average nucleosome occupancy relative to the TSS (in bp) of (A) all protein-coding genes, (B) expressed protein-coding genes, (C) unexpressed protein-coding genes, (D) genes with expression changes in rH4R17A, *arid5, rlt1/2,* and/or *chr11/17* mutants, (E) genes with no expression changes in rH4R17A, *arid5, rlt1/2,* and/or *chr11/17* mutants, (F) upregulated genes and (G) downregulated genes in rH4R17A, *arid5, rlt1/2,* and/or *chr11/17* mutants. The MNase-seq results were generated from two independent biological replicates and RNA-seq data were obtained from the same tissues used for MNase-seq. Cutoffs were defined as follows: Expressed ≥0.5 TPM; Unexpressed <0.5 TPM. Genes with expression changes were defined as >±1.5-fold vs. Col and genes with no expression changes were defined as <±1.1-fold vs. Col. (H-N) Fold change in ΔNucleosome Occupancy of +2 through +6 nucleosome peaks relative to Col corresponding to (H) all protein-coding genes, (I) expressed protein-coding genes, (J) unexpressed protein-coding genes, (K) genes with expression changes in rH4R17A, *arid5, rlt1/2,* and/or *chr11/17* mutants, (L) genes with no expression changes in rH4R17A, *arid5, rlt1/2,* and/or *chr11/17* mutants, (M) upregulated genes and (N) downregulated genes in rH4R17A, *arid5, rlt1/2,* and/or *chr11/17* mutants. Standard deviation denoted with error bars. *P*-value from paired Student’s *t*-test denoted with asterisks (*p<0.05, **p<0.005, ***p<0.0005).

To further investigate the interplay in plants between H4R17 and ISWI, we performed whole-transcriptome analysis via RNA sequencing (RNA-seq) on the rH4R17A, *arid5, rlt1/*2, *chr11/17* and *pie1* (catalytic subunit of the SWR1 complex) mutants grown in short-day conditions. Our results from the RNA-seq analyses showed that there were 1771 downregulated genes and 1471 upregulated genes in *chr11/17* double mutants (3242 differently expressed genes [DEGs] in total), while there were only 535 downregulated genes and 299 upregulated genes in the rH4R17A-1 mutant (834 DEGs), and 410 downregulated genes and 375 upregulated genes in the rH4R17A-2 mutant (785 DEGs) (Fig. 4E). In spite of the large difference in the total amount of DEGs between *chr11/17* mutants and the rH4R17A plants, we observed a high overlap between the DEGs in the *chr11/17* and rH4R17A-1 mutants (45.6%, 380/834), as well as the DEGs in the *chr11/17* and rH4R17A-2 mutants (56.0%, 440/785) (Fig. 4E). Additionally, we observed a high overlap of DEGs (average 50.0% rH4R17A vs. *arid5*; average 52.5% rH4R17A vs. *rlt1/2*) when comparing the rH4R17A lines with the ISWI subunit mutants *arid5* and *rlt1/2* (Fig. 4F-G). Furthermore, in the rH4R17A, *chr11/17*, *arid5*, and *rlt1/2* mutants, we detected a similar pattern of RNA expression not shared by rH4 or Col plants, indicated by the clustering of both rH4R17A lines and all ISWI subunit mutants together (Fig. 4H). In contrast, we did not observe substantial overlap (average 25.8%) between the DEGs identified in *pie1* mutants and the DEGs of rH4R17A mutants (Fig. 4H, Supp. Fig. 6A). We then investigated the expression of key flowering time regulatory genes and found that the flowering promoter genes *FRUITFULL* (*FUL*), *SOC1* and *FT* were all co-upregulated in the rH4R17A, *rlt1/2*, *arid5,* and *chr11/17* mutants (Fig. 4I, Supp. Fig. 6B). These patterns of co-expression were not observed when comparing *rlt1/2*, *arid5,* and *chr11/17* mutants to rH4 plants (Fig. 4H-I, Supp. Fig. 6B). The shared developmental phenotypes and transcriptional profiles of the rH4R17A, *rlt1/2*, *arid5,* and *chr11/17* mutants suggest that H4R17 plays an important role in plants as in other eukaryotes in regulating the activity of ISWI on chromatin.

**Figure 6:**
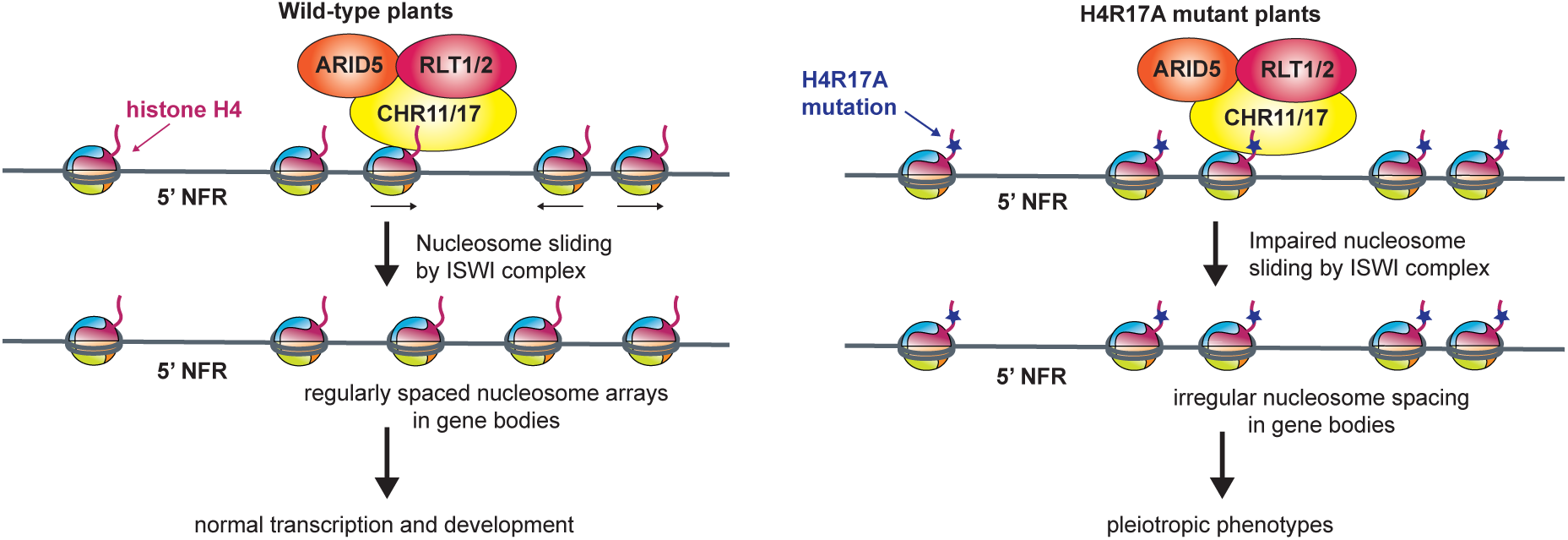
Model for the role of H4R17 in plants. Proposed model for the role of histone H4 arginine 17 in the regulation of ISWI complexes in *A. thaliana*. 5’ NFR: 5’ Nucleosome-Free Region.

### Impact of H4R17A mutation on global nucleosome positioning

ISWI functions as a chromatin remodeling complex that properly organizes nucleosome spacing at transcriptionally active genes in eukaryotes (Clapier and Cairns, 2009; Gkikopoulos et al., 2011; Li et al., 2014; Yadon and Tsukiyama, 2011). Due to the similarity in the phenotypes and transcriptional profiles between rH4R17A plants and mutants in the *Arabidopsis* ISWI complex, we hypothesized that expression of H4R17A interferes with nucleosome spacing in plants. To address this hypothesis, we assessed global nucleosome positioning in rH4R17A mutants using micrococcal nuclease digestion followed by deep sequencing (MNase-seq). Consistent with previous results, a relatively lower nucleosome density was found in the 1-kb region upstream of the transcription start site (TSS) of protein-coding genes, while a relatively high density, evenly spaced nucleosome distribution was found in the 1-kb region downstream of the TSS for Col plants (Fig. 5A) (Li et al., 2014). Moreover, expressed protein-coding genes were generally observed to display more highly phased nucleosome arrays in the gene body and a sharper peak of nucleosome-free DNA in the promoter when compared to unexpressed protein-coding genes, in line with previous studies (Fig. 5B-C) (Li et al., 2014; Zhang et al., 2015). In terms of the different genotypes analyzed, rH4 plants displayed highly similar nucleosome positioning patterns to Col as expected. In contrast, while rH4R17A, *arid5*, and *rlt1/2* mutants displayed the same general pattern of lower nucleosome density upstream of the TSS and high nucleosome density downstream of the TSS, these genotypes all exhibited a reduction of evenly spaced nucleosome distributions in the gene body (Fig. 5A), similar to the pattern reported for the *chr11/17* mutant (Li et al., 2014). Additionally, we analyzed the nucleosome distribution patterns at genes with expression changes in rH4R17A, *chr11/17*, *rlt1/2*, and/or *arid5* mutants as well as genes without expression changes in these mutants. We found that the nucleosome distribution patterns at DEGs and non-DEGs were both affected by the rH4R17A, *arid5,* and *rlt1/2* mutations (Fig. 5D-E), in line with previously published MNase-seq results for the *chr11/17* mutant (Li et al., 2014). Additionally, nucleosome distribution patterns at DEGs were affected by the rH4R17A, *arid5,* and *rlt1/2* mutations regardless of whether the expression of these genes was up- or downregulated (Fig. 5F-G). To provide a more quantitative assessment of nucleosome spacing in our assays, we calculated the average change in nucleosome occupancy at the +2 through +6 nucleosome peaks as a measure of nucleosome phasing. This analysis confirmed that the rH4R17A, *arid5,* and *rlt1/2* mutations caused a significant reduction in regular nucleosome phasing in gene bodies (Fig. 5H-N). Taken together, these results indicate that H4R17 positively regulates the action of the ISWI complex to establish nucleosome arrays in protein-coding genes.

## Discussion

### A novel system for studying histone function in plants

In this study, we present a new histone replacement system that facilitates the analysis of histone H4 functions on diverse processes in plants. Our results serve as a proof-of-concept that complete histone replacement systems can be rapidly established in *A. thaliana*. In the future, this approach may be applied to generate similar systems to study the functions of different histones or histone variants. The histone replacement system developed in this study for histone H4 will supplement already existing systems in *S. cerevisiae* and *D. melanogaster* to offer new biological insights into the roles of H4 in plants. In its current iteration, our methodology already provides the most extensive coverage of H4 mutants in a multicellular eukaryote, as the histone replacement system generated in *D. melanogaster* has only been used to generate 14 H4 point mutants (Zhang et al., 2019), compared to the 63 H4 point mutants generated with our system, which will be made available through the Arabidopsis Biological Resource Center (ABRC).

Our CRISPR-based strategy to replace endogenous histones offers several advantages over other methods that can potentially be used to achieve complete histone replacement. For example, the successful generation of the H4 septuple mutant in two generations in this work demonstrates that multiplex CRISPR/Cas9 can be used to efficiently inactivate a large number of histone genes in plants (LeBlanc et al., 2017). Using CRISPR/Cas9 greatly reduces the amount of time and resources required to generate a histone depletion background, especially when compared to crossing individual histone mutants. The presence of tandem duplicated copies of histone genes (e.g. the H3.1 genes *At5g10390* and *At5g10400*) can also preclude using traditional crossing schemes to generate backgrounds lacking a specific histone or histone variant. In addition, deploying multiplex CRISPR/Cas9 to inactivate endogenous histones will allow researchers to rapidly re-establish histone replacement systems in a particular mutant background, for example, to screen for point mutations in histones that enhance or suppress a phenotype of interest. Another advantage of our histone H4 replacement strategy is that we consistently observed high expression of the H4 replacement gene, which rescues the morphological phenotype of the H4 septuple background in all of our T1 lines (Fig. 1A-C). Several factors could contribute to this phenomenon. While T-DNA integration into the *A. thaliana* genome occurs randomly, selection pressure appears to shift the recovery of T-DNA insertions into more transcriptionally active chromatin regions (Alonso et al., 2003; Brunaud et al., 2002; Kim et al., 2007; Koncz et al., 1989; Szabados et al., 2002). Moreover, dosage compensation mechanisms acting to up-regulate the expression of the endogenous histone H4 gene *At3g53730,* as seen in this study (Fig. 1C), may also act on the histone H4 replacement gene, as the expression of both of these genes is driven by the native *At3g53730* promoter. Histone dosage compensation has also been observed in the histone replacement systems implemented in the multicellular eukaryote *D. melanogaster* (McKay et al., 2015; Zhang et al., 2019). These histone dosage compensation mechanisms may be related to the recently described process of transcriptional adaptation, in which mutant mRNA decay causes the upregulation of related genes (El-Brolosy et al., 2019; Serobyan et al., 2020). For both of the above reasons, transgenic plants lacking endogenous H4 proteins are observed, and therefore, rH4 plants expressing exclusively mutant histones can predictably be obtained using our strategy.

Several changes could be implemented to improve future histone replacement systems in *A. thaliana* and other plants. To control for differential effects caused by random T-DNA integration (Gelvin, 2017), we characterized in this study two independent transgenic lines expressing each H4 replacement construct. Ideally, gene targeting would be utilized to introduce the H4 mutations directly at an endogenous histone H4 locus. While gene targeting technologies in plants currently have very low efficiency compared to yeast and animal systems, as additional improvements in gene targeting are developed, *in situ* histone replacement systems in plants analogous to platforms already existing in yeast and *Drosophila melanogaster* may also become feasible. Additionally, more precise control over the dosage of the replacement histone could also serve to improve this method. It was recently shown that while yeast histone replacement systems utilizing single-copy integrated histone genes expressing certain mutant histones cannot survive, the addition of a second copy of the mutant histone gene rescues this lethality (Jiang et al., 2017). In the system described here, we utilized a single endogenous histone H4 gene for the H4 replacement gene, rather than generating eight histone H4 replacement genes corresponding to each endogenous histone H4 gene present in the *A. thaliana* genome. While we observed that our rH4 plants appear morphologically wild-type due to high H4 expression, it may be important to study the function of the other endogenous H4 genes, or the requirement for *A. thaliana* to have eight copies of the H4 genes in its genome. Although labor-intensive, future strategies simultaneously using multiple endogenous H4 genes as H4 replacement genes could therefore be more reflective of the H4 supply available to wild-type plants.

### H4R17 regulates nucleosome remodeling and developmental processes in plants

This study has identified a novel role for H4R17 in regulating multiple developmental processes in *A. thaliana*, including leaf development, fruit development and flowering. Our findings suggest that this role for H4R17 is not mediated via post-translational modification of this residue. Based on our results, we propose a model similar to that of animal systems where H4R17 regulates developmental processes in plants through its regulation of the ISWI complex (Fig. 6) (Clapier et al., 2001; Clapier et al., 2002; Dann et al., 2017; Fazzio et al., 2005; Hamiche et al., 2001). In wild-type plants, H4R17 positively regulates the ISWI complex to slide nucleosomes and adequately establish the nucleosome positioning patterns in the gene bodies of protein-coding genes (Fig. 6). In rH4R17A mutant plants however, the positive regulation of ISWI by histone H4 is impaired so that evenly-spaced nucleosome distributions are no longer observed in gene bodies. The altered nucleosome positioning patterns in gene bodies and the large-scale transcriptional changes in turn cause the observed pleiotropic developmental phenotypes. Comparative analysis of the protein sequence of the ISWI catalytic subunits in *A. thaliana* (CHR11 and CHR17) reveals strict conservation of the amino acids involved in making contacts with histone H4 arginine 17 in the ISWI orthologs from other species (Supp. Fig. 10) (Yan et al., 2016; Yan et al., 2019), which supports the findings of this study. Additionally, homology modeling of CHR11 indicates structural conservation of the H4R17-binding region in plant ISWI proteins (Supp. Fig. 11). Interestingly, while transcription and nucleosome positioning have been shown to be highly interconnected processes (Hughes et al., 2012; Jiang and Pugh, 2009; Struhl and Segal, 2013; Workman and Kingston, 1998), we and others have observed that mutations in H4R17 and plant ISWI complex subunits affect nucleosome positioning patterns in both differentially and non-differentially expressed genes (Li et al., 2014). Our results support previous work demonstrating that it is unlikely that the nucleosome positioning defects in ISWI mutants are caused by the transcriptional changes observed in these backgrounds (Li et al., 2014; Luo et al., 2020). Moreover, our results are consistent with the idea that many factors on top of nucleosome positioning in gene bodies affect the transcription level of a gene, and thus in some cases, altered genic nucleosome positioning appears to majorly impact transcription, while in others, little change is observed (Bai and Morozov, 2010; Jiang and Pugh, 2009). Additionally, processes related to genetic robustness may also serve to counteract transcriptional fluctuations due to perturbations of nucleosome positioning (Masel and Siegal, 2009). For these reasons, we observed independence between the nucleosome positioning and transcriptional phenotypes of rH4R17A and ISWI mutants.

ISWI chromatin remodeling complexes contain between two and four subunits in eukaryotes, including a conserved ATPase catalytic subunit and at least one accessory subunit (Aydin et al., 2014; Clapier and Cairns, 2009; Corona and Tamkun, 2004). Multiple types of ISWI complexes have been identified in animals, and the different accessory subunits in these complexes have been proposed to modulate the activity of the shared catalytic subunit as well as the specificity and target recognition of the complex (Aydin et al., 2014; Lusser et al., 2005; Toto et al., 2014). In plants, there are three types of ISWI complexes that have been identified: the plant-specific CHR11/17-RLT1/2-ARID5 (CRA)-type complex, the CHR11/17-DDP1/2/3-MSI3 (CDM)-type complex, and the CHR11/17-DDR1/3/4/5-DDW1 (CDD)-type complex (Tan et al., 2020). In addition, the shared ISWI catalytic subunits CHR11 and CHR17 were also recently demonstrated to act as accessory subunits of the SWR1 chromatin remodeling complex in plants (Luo et al., 2020). Given that there are multiple types of ISWI complexes in plants, we demonstrated that mutations in the CRA-type complex demonstrate a less severe impact on nucleosome positioning than rH4R17A mutations, which is in line with the model that H4R17 regulates all three types of ISWI complexes through its interaction with the catalytic subunits CHR11 and CHR17 (Clapier et al., 2001; Clapier et al., 2002; Dann et al., 2017; Fazzio et al., 2005; Hamiche et al., 2001). Further characterization of the different ISWI complexes in plants, including their different targeting specificities to chromatin loci and the impact of the other identified CDM-type and CDD-type complexes on the regulation of global transcription and nucleosome positioning, will contribute to elucidating their specific consequences on chromatin regulation.

## Materials and Methods

### Plant materials

All *Arabidopsis* plants were derived from the Columbia ecotype and grown in Pro-Mix BX Mycorrhizae soil under cool-white fluorescent lights (approximately 100 μmol m^−2^ s^−1^). Seeds were surface-sterilized with a 70% ethanol, 0.1% Triton solution for 5 minutes, and then with 95% ethanol for one minute. Seeds were spread on sterilized paper and plated on 0.5% Murashige-Skoog (MS) plates. Seeds were stratified in the dark at 4°C for 2 to 4 days, transferred to the growth chamber for 5 days, and then transplanted to soil. Plants grown in long-day conditions were grown for 16 h light/ 8 h dark, and plants grown in short-day conditions were grown for 8 h light/ 16 h dark.

The *chr11* (GK-424F01)/ *chr17* (GK-424F04) double mutant was described previously (Li et al., 2012). The *arid5* (SALK_111627), *prmt7-1* (SALK_028160), and *prmt7-2* (SALK_039529) T-DNA insertion mutants were obtained from the Arabidopsis Biological Resource Center. The *pie1* T-DNA insertion mutants were initially obtained from the Arabidopsis Biological Resource Center (SALK_096434) and the *pie1* mutants used in this study were seeds collected from homozygous *pie1* plants. The *rlt1* (SALK_099250)/ *rlt2* (SALK_132828) double mutants were generated by crossing. Due to severely reduced fertility, *chr11*/*17* and *arid5* mutants were maintained in a heterozygous state.

### Generation of transgenic *Arabidopsis* plants

Binary vectors were transformed into *Agrobacterium tumefaciens* (strain GV3101) using heat shock and plants were transformed with these constructs using the floral dip procedure as described previously (Clough and Bent, 1998). Transgenic plants for generation of the H4 septuple mutant were selected on 0.5 MS plates containing 1% sucrose, carbenicillin (200 μg ml^−1^) and kanamycin (100 μg ml^−1^). Transgenic rH4 plants were selected on 0.5 MS plates containing 1% sucrose, carbenicillin (200 μg ml^−1^) and glufosinate ammonium (25 μg ml^−1^). Plants were subjected to heat stress treatments as described previously (LeBlanc et al., 2017). The plants were grown continuously at 22°C from that point on.

### Flowering time and rosette measurements

Days to flower was measured when a 1 cm bolting stem was visible. Rosette leaves were counted at day of flowering. Rosette area was measured using the ARADEEPOPSIS workflow (Huther et al., 2020).

### Dimensionality reduction and clustering

Principal component analysis of four variables (day number in long-day, leaf number in long-day, day number in short-day, and leaf number in short-day) was performed. We centered variables at mean 0 and set the standard deviation to 1. *k*-means clustering was performed 40 times with random initializations on the first two principal components to identify three clusters. Analyses were performed in RStudio with R version 3.6.1 (Team, 2018).

### Plasmid construction

CRISPR constructs used to generate the H4 septuple mutant were inserted into the *pYAO*-Cas9-SK vector as described previously (Yan et al., 2015).

The H4 replacement plasmid was made by amplifying the promoter (967 bp upstream of start codon), gene body, and terminator (503 bp downstream of stop codon) of H4 (*At3g53730*) into pENTR/D (ThermoFisher Scientific, Waltham, MA, USA). Site-directed mutagenesis of *pH4::H4* in pENTR/D using QuikChange II XL (Agilent Technologies, Santa Clara, CA, USA) was first performed to create plasmids with 10 silent mutations in the H4 coding sequence. These silent mutations were engineered to test the resistance of the H4 replacement gene against multiple gRNAs. Additional site-directed mutagenesis of this vector was performed to generate a library of 63 H4 point mutant genes.

Each *pH4*::*H4* sequence was then transferred into the binary vector pB7WG, containing the H4 gRNA, using Gateway Technology. The binary vector pB7WG containing the H4 gRNA was generated as follows: The AtU6-26-gRNA vector containing the gRNA targeting H4 (*At3g53730*) was first digested with the restriction enzymes *SpeI* and *NheI*, and the digestion products were run on a 1% agarose gel. Then, the band containing the H4 gRNA was cut out and ligated into the binary vector pB7WG, which had been digested with the restriction enzyme *SpeI*.

The *PRMT7* overexpression construct was created by cloning the genomic *PRMT7* gene (from ATG to stop codon, including introns) into pDONR207, and then subcloning the gene into pMDC32 (Curtis and Grossniklaus, 2003).

### DNA extraction, PCR and sequencing analyses

Genomic DNA was extracted from *Arabidopsis* plants by grinding one leaf in 500 μl of Extraction Buffer (200 mM of Tris-HCl pH8.0, 250 mM NaCl, 25 mM ethylenediaminetetraacetic acid (EDTA) and 1% SDS). Phenol/chloroform (50 μl) was added and tubes were vortexed, followed by centrifugation for 10 min at 3220*g*. The supernatant was transferred to a new tube and 70 μl of isopropanol was added, followed by centrifugation for 10 min at 3200*g*. The supernatant was removed and the DNA pellets were resuspended in 100 μl of water.

PCR products were sequenced and analyzed using Sequencher 5.4.6 (Gene Codes Corporation, Ann Arbor, MI, United States) to identify CRISPR-induced mutations. To assess the rate of mutation of the remaining endogenous H4 gene (*At3g53730*) in rH4 plants by the gRNA in the H4 replacement plasmid, endogenous H4 PCR products were cloned into TOPO TA cloning vectors (Invitrogen, Carlsbad, CA, United States). Ten to sixteen individual clones corresponding to each plant were sequenced.

### RT-qPCR

RNA was extracted from 4-week-old leaf tissue with TRIzol (Invitrogen) and DNase treated using RQ1 RNase-Free DNase (Promega, Madison, WI, USA). Three biological replicates (different plants sampled simultaneously) were assessed. SuperScript II Reverse Transcriptase (Invitrogen) was used to produce cDNA. Reverse transcription was initiated using random hexamers (Applied Biosystems, Foster City, CA, United States). Quantification of cDNA was done by real-time PCR using a CFX96 Real-Time PCR Detection System (Bio-Rad, Hercules, CA, USA) with KAPA SYBR FAST qPCR Master Mix (2x) Kit (Kapa Biosystems, Wilmington, MA, USA). Relative quantities were determined by using a comparative Ct method as follows: Relative quantity = 2(−((Ct GOI unknown − Ct normalizer unknown) − (Ct GOI calibrator − Ct normalizer calibrator))), where GOI is the gene of interest (Livak and Schmittgen, 2001). Actin was used as the normalizer.

### Next-generation sequencing library preparation

RNA-seq and MNase-seq libraries were prepared at the Yale Center for Genome Analysis (YCGA). Leaves of 4-week-old plants grown in short-day conditions were frozen in liquid nitrogen, ground with a mortar and pestle, and then RNA was extracted using the RNeasy Plant Mini Kit (Qiagen, Hilden, Germany). RNA quality was confirmed through analysis of Agilent Bioanalyzer 2100 electropherograms (Supp. Fig. 7). Library preparation was performed using Illumina’s TruSeq Stranded Total RNA with Ribo-Zero Plant in which samples were normalized with a total RNA input of 1 μg and library amplification with 8 PCR cycles. MNase-digested DNA was collected as described previously (Pajoro et al., 2018) with the following modifications: 2 g of leaf tissue from 4-week-old plants grown in short-day conditions was ground in liquid nitrogen and resuspended in 20 ml of lysis buffer for 15 minutes at 4°C. The resulting slurry was filtered through a 40 μm cell strainer into a 50 ml tube. Samples were centrifuged for 20 minutes at 3200*g*. The resulting pellets were resuspended in 10 ml of HBB buffer and centrifuged for 10 minutes at 1500*g*. Pellets were successively washed in 5 ml wash buffer and 5 ml reaction buffer. MNase-seq library preparation was performed using the KAPA Hyper Library Preparation kit (KAPA Biosystems, Part#KK8504). For each biological replicate, pooled leaf tissue collected simultaneously from three different plants was used. Libraries were validated using Agilent Bioanalyzer 2100 Hisense DNA assay and quantified using the KAPA Library Quantification Kit for Illumina® Platforms kit. Sequencing was done on an Illumina NovaSeq 6000 using the S4 XP workflow.

### RNA-seq processing and analysis

Two independent biological replicates for Col, rH4-1, rH4-2, rH4R17A-1, rH4R17A-2, *arid5, rlt1/2, chr11/17,* and *pie1* were sequenced. Paired-end reads were filtered and trimmed using Trim Galore! (version 0.5.0) with default options for quality (https://github.com/FelixKrueger/TrimGalore). The resulting data sets were aligned to the Araport11 genome (Cheng et al., 2017) using STAR (version 2.7.2a) allowing 2 mismatches (-- outFilterMismatchNmax 2) (Dobin et al., 2013). Statistics for mapping and coverage of the RNA-seq data are provided in Supplemental Data Set 1. Protein-coding genes were defined as described in the Araport11 genome annotation (Cheng et al., 2017). The program featureCounts (version 1.6.4) (-M --fraction) (Liao et al., 2014) was used to count the paired-end fragments overlapping with the annotated protein-coding genes. Differential expression analysis of protein-coding genes was performed using DESeq2 version 1.26 (Love et al., 2014) on raw read counts to obtain normalized fold changes and *Padj*-values for each gene. Genes were considered to be differentially expressed if they showed >±2-fold-change and *Padj*-value <0.05. Differentially expressed genes are described in Supplemental Data Set 2. Venn diagrams, correlation plots and correlation matrices were plotted using RStudio with R version 3.6.1 (Team, 2018). Heatmaps were plotted with the pheatmap package (version 1.0.12) in RStudio using default clustering parameters on rows and columns. Consistency between biological replicates was confirmed by Spearman correlation using deepTools2 (version 2.7.15) (Supp. Fig. 8) (Ramirez et al., 2016). deepTools2 was used to generate bam coverage profiles for visualization with Integrative Genomics Viewer version 2.8.9 (Robinson et al., 2011).

### MNase-seq processing and analysis

Two independent biological replicates for Col, rH4-2, rH4R17A-1, *arid5*, and *rlt1/2* were sequenced. Paired-end reads were filtered and trimmed using Trim Galore! (version 0.5.0) with default options for quality (https://github.com/FelixKrueger/TrimGalore). Bowtie2 version 2.4.2 (Langmead and Salzberg, 2012) was used to align the reads to the Araport11 genome (Cheng et al., 2017) with the --very-sensitive parameter. Statistics for mapping and coverage of the MNase-seq data are provided in Supplemental Data Set 1. Duplicate reads were removed using Picard toolkit version 2.9.0 (toolkit., 2019) (MarkDuplicates with *REMOVE_DUPLICATES=true*) and the insertion size was filtered from 140 bp to 160 bp using SAMtools version 1.11 (Li et al., 2009). The average nucleosome occupancy corresponding to the regions 1-kb upstream and downstream of the TSS of all protein-coding genes was calculated using the bamCoverage (--MNase parameter specified) and computeMatrix functions of deepTools2 version 2.7.15 (Ramirez et al., 2016). Normalization was performed by scaling with the effective library size calculated by the calcNormFactors function using edgeR version 3.28.1 (Robinson et al., 2010). Consistency between biological replicates was confirmed by Spearman correlation using deepTools2 (Supp. Fig. 9). Fold change in ΔNucleosome Occupancy of +2 through +6 nucleosome peaks relative to Col was calculated with a custom Python script (https://github.com/etc27/MNaseseq-workflow/analysis/peak_height) as follows: ΔNucleosome Occupancy = peak maximum – (5’ peak minimum + 3’ peak minimum)/2.

### Model building

The homology model for *Arabidopsis thaliana* CHR11 (a.a. 176-706) was built with Swiss-Model against the *Myceliophthora thermophila* ISWI reference structure (5JXR) (Biasini et al., 2014; Yan et al., 2016).

### Primers

All primers used in this study are listed in Supplemental Data Set 3 (Dong et al., 2021; Richter et al., 2019; Wu et al., 2008).

### Statistical analyses

Statistical analysis data are provided in Supplemental Data Set 4.

## Data Availability Statement

Raw and processed RNA-seq and MNase-seq data have been deposited in the Gene Expression Omnibus database with the accession code GSE190317.

## Accession Numbers

Accession numbers of genes reported in this study include: AT3G53730 (*H4*), AT1G07660 (*H4*), AT1G07820 (*H4*), AT2G28740 (*H4*), AT3G45930 (*H4*), AT5G59690 (*H4*), AT3G46320 (*H4*), AT5G59970 (*H4*), AT4G21070 (*BRCA1*), AT1G65480 (*FT*), AT2G45660 (*SOC1*), AT4G16570 (*PRMT7*), AT3G06400 (*CHR11*), AT5G18620 (*CHR17*), AT3G43240 (*ARID5*), AT1G28420 (*RLT1*), AT5G44180 (*RLT2*), AT3G12810 (*PIE1*), AT5G60910 (*FUL*), and AT5G09810 (*ACTIN7*)

## Supplemental Data

Supplemental Data Set 1: Statistics for mapping and coverage of the NGS data.

Supplemental Data Set 2: Differentially Expressed Genes identified in RNA-seq.

Supplemental Data Set 3. Cloning and PCR primers.

Supplemental Data Set 4. Statistical analysis data.

## Acknowledgments

We thank all current and former members of our lab for discussions and advice during the course of this work. We especially acknowledge the contributions of Anisa Iqbal, Benoit Mermaz and Gonzalo Villarino. We acknowledge Christopher Bolick and his staff at Yale for help with plant growth and maintenance and we thank the Yale Science Building Facilities Staff for maintenance of the lab facilities. We thank Franziska Bleichert from Yale University for her assistance with structural analyses. This project was supported by grant #R35GM128661 from the National Institutes of Health to Y.J. E.T.C. was supported by a Yale University Gruber Science Fellowship, the NIH Predoctoral Program in Cellular and Molecular Biology Training Grant T32GM007233, and the National Science Foundation Graduate Research Fellowship #2139841. U.V.P. was supported by a grant from National Institutes of Health R35GM125003. We thank Paja Sijacic and Roger Deal from Emory University for their generous gift of *pie1* seeds. We thank Lin Xu and Wu Liu from the National Laboratory of Plant Molecular Genetics, Shanghai Institute of Plant Physiology and Ecology, Shanghai Institutes for Biological Sciences, Chinese Academy of Sciences for their generous gift of *chr11/17* mutant seeds. The authors declare that they have no competing interests.

## Author Contributions

Y.J. and E.T.C. designed the experiments. E.T.C. and Y.J. wrote the paper with contributions from C.L. Constructs were generated by E.T.C. Plant transformations were performed by E.T.C., A.S., C.L, and Y.J. Genotyping was performed by E.T.C., A.S., M.A.T, C.L, and Y.J. RNA extractions and RT-qPCR were done by E.T.C., C.L. and Y.J. Flowering time measurements were obtained by E.T.C., M.A.T. and C.L. Plant pictures were taken by E.T.C., M.A.T., C.L. and Y.J. Some of the mutants used in this work were generated by Y.H. and U.V.P. C.L. performed the RNA-seq and MNase-seq experiments. E.T.C. did the bioinformatics analyses of all RNA-seq and MNase-seq experiments.

## Supplemental Figures

**Supplemental Figure 1:**
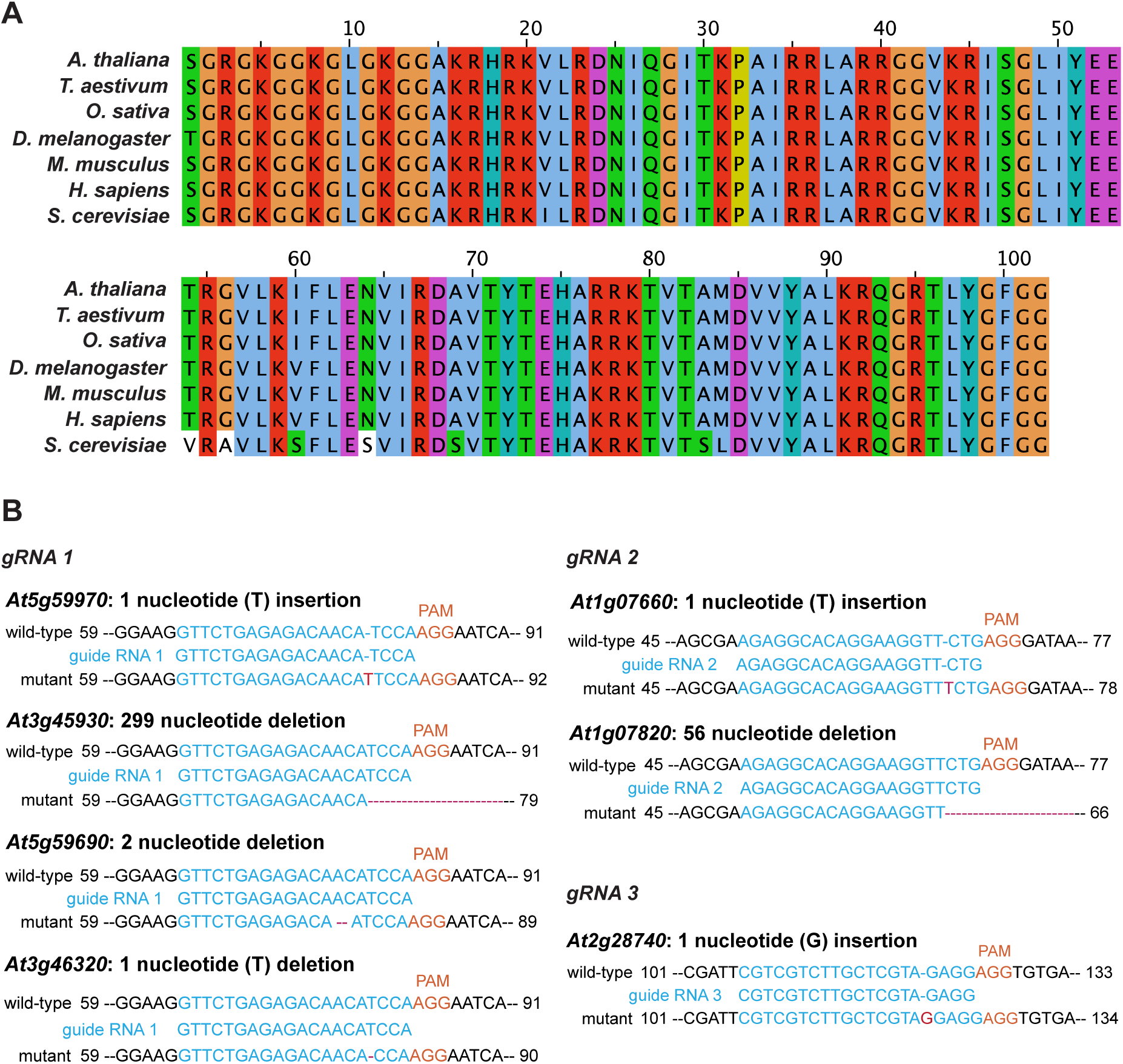
CRISPR/Cas9-induced mutations in the H4 septuple mutant of *A. thaliana*. (A) Multiple sequence alignment of histone H4 proteins performed with Clustal Omega. Protein sequences were obtained from UniProt and correspond to the following accession numbers: *Arabidopsis thaliana*; P59259, *Triticum aestivum*; P62785*, Oryza sativa*; Q7XUC9*, Drosophila melanogaster*; P84040*, Mus musculus*; P62806*, Homo sapiens*; P62805, and *Saccharomyces cerevisiae*; P02309. Chemical characteristics of amino acids shown with ClustalX color scheme (Larkin et al., 2007). (B) Design of the three gRNAs targeting seven endogenous histone H4 genes in *A. thaliana.* The resulting homozygous mutation in each of the targeted genes in the H4 septuple mutant is shown.

**Supplemental Figure 2:**
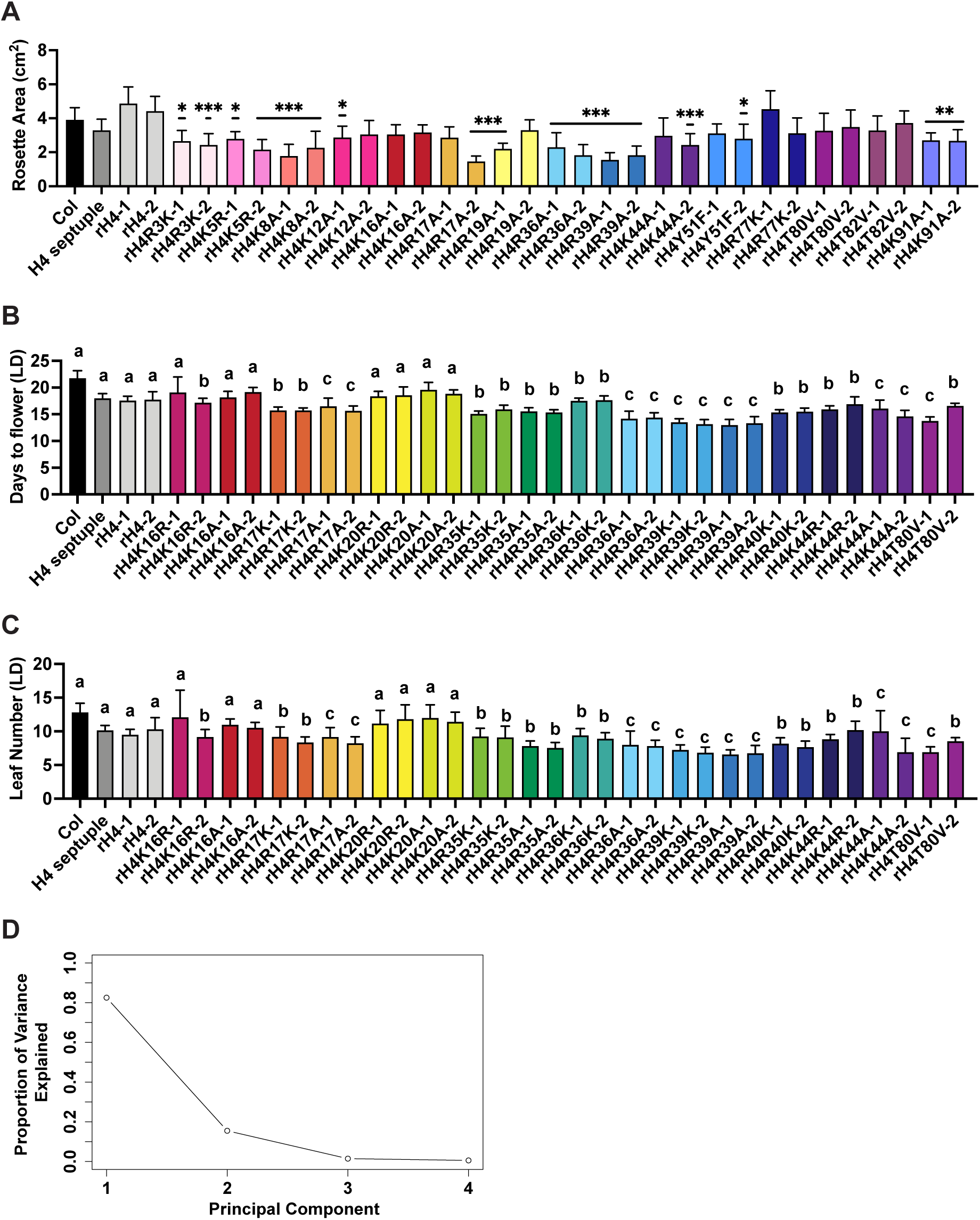
Impact of histone H4 mutations on the floral transition in *A. thaliana*. (A) Mean rosette area of Col, the H4 septuple mutant, and various H4 replacement backgrounds (two independent transgenic lines each). Standard deviation shown with error bars (n≥9). Statistical analyses were performed using one-way ANOVA with Tukey’s HSD post hoc test. *P*-value from Tukey’s HSD test (genotype vs. Col) denoted with asterisks (*p<0.05, **p<0.005, ***p<0.0005). (B-C) Mean (B) days to flower and (C) rosette leaf number at flowering in long-day (LD) conditions for Col, the H4 septuple mutant, and various H4 replacement backgrounds (two independent transgenic lines each). Standard deviation shown with error bars (n≥11). Letters (a,b,c) indicate cluster identified by *k*-means clustering. (D) Scree plot depicting the proportion of variance explained by each of the principal components.

**Supplemental Figure 3:**
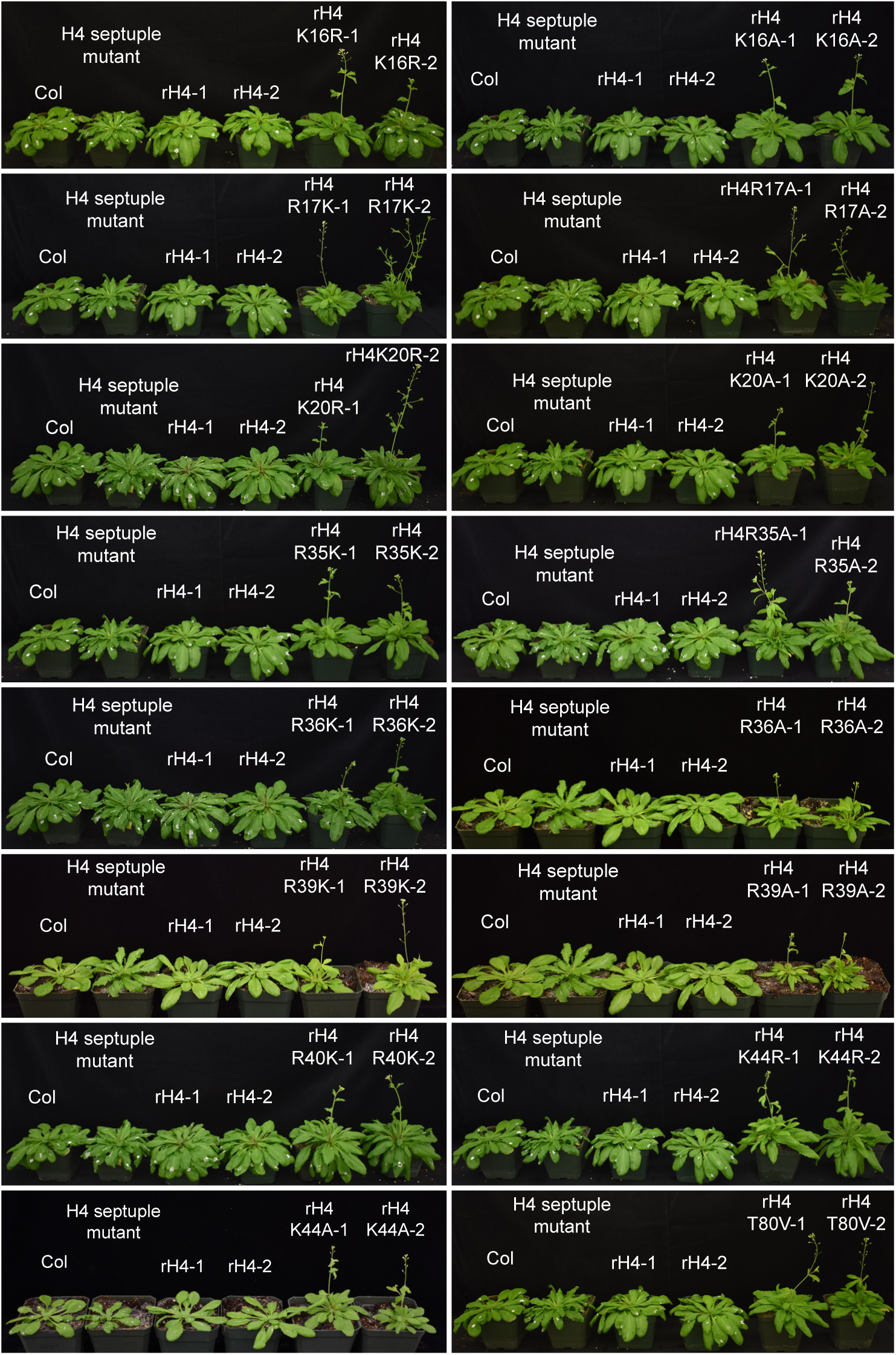
Phenotypes of early flowering histone H4 mutants. Phenotype of Col, the H4 septuple mutant, and various H4 replacement backgrounds grown in short-day between 5 and 7.5 weeks. Two independent transgenic lines were assessed per H4 point mutation. White marks present on certain rosette leaves due to leaf counting measurements.

**Supplemental Figure 4:**
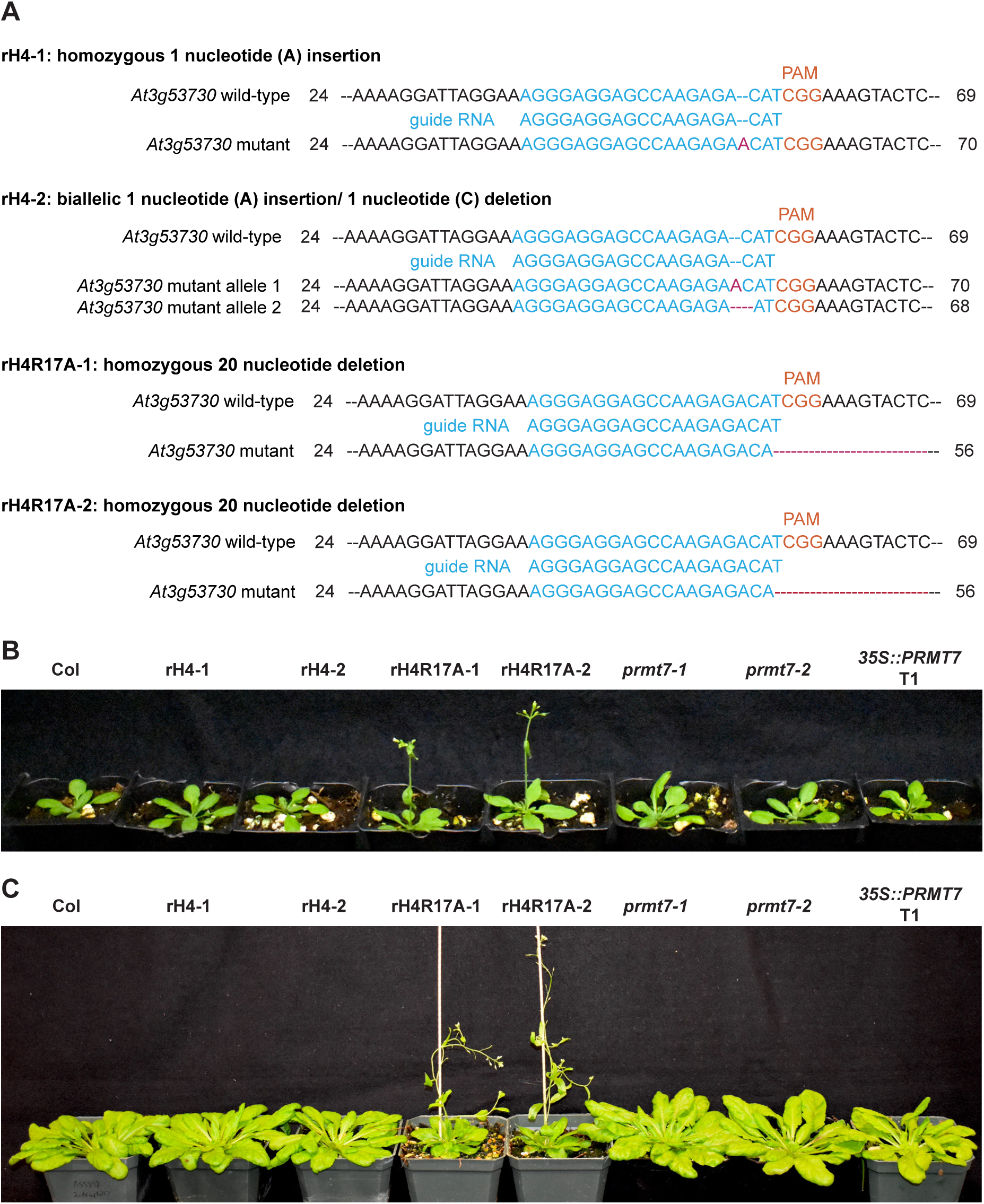
No functional relationship between rH4R17A and *PRMT7* mutations. (A) Homozygous or biallelic mutation in histone H4 (*At3g53730*) in rH4-1, rH4-2, rH4R17A-1 and rH4R17A-2 plants. (B-C) Phenotype of Col, rH4-1, rH4-2, rH4R17A-1, rH4R17A-2, *prmt7-1, prmt7-2*, and *35S::PRMT7* T1 plants grown in (B) long-day at 3 weeks and (C) short-day at 8 weeks.

**Supplemental Figure 5:**
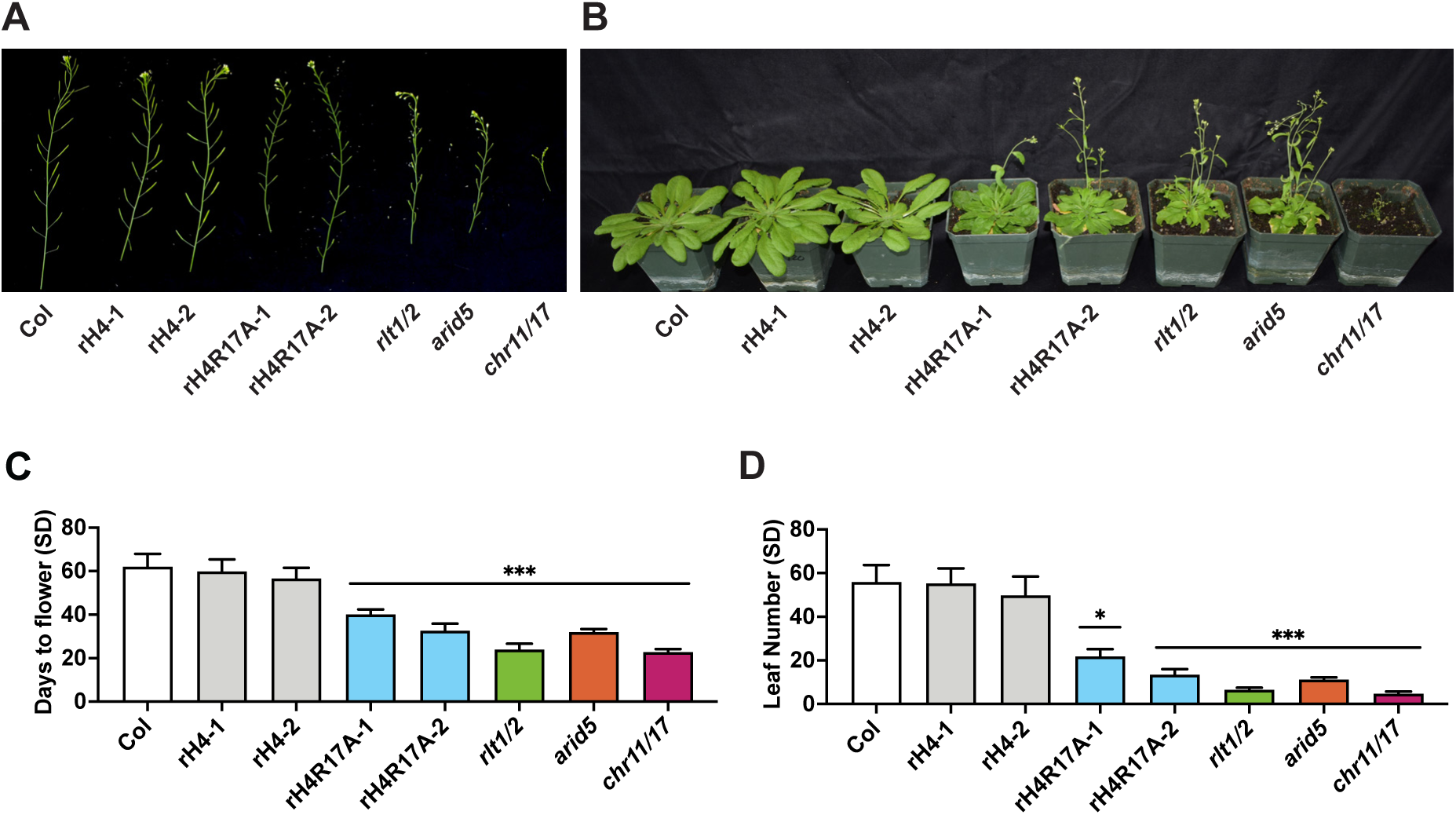
The effect of ISWI and rH4R17A mutations on the floral transition and development. (A) Siliques of Col, rH4-1, rH4-2, rH4R17A-1, rH4R17A-2, *rlt1/2, arid5*, and *chr11/17* plants grown in long-day for 4 weeks. (B) Morphological phenotypes of Col, rH4-1, rH4-2, rH4R17A-1, rH4R17A-2, *rlt1/2, arid5*, and *chr11/17* plants grown in short-day at 7 weeks. (C-D) Mean (C) days to flower and (D) rosette leaf number at flowering in short-day (SD) conditions for Col, rH4-1, rH4-2, rH4R17A-1, rH4R17A-2, *rlt1/2, arid5*, and *chr11/17* plants. Standard deviation shown with error bars. Statistical analyses were performed using one-way ANOVA with Tukey’s HSD post hoc test. *P*-value from Tukey’s HSD test (genotype vs. Col) denoted with asterisks (*p<0.01, **p<0.001, ***p<0.0001). n=12.

**Supplemental Figure 6:**
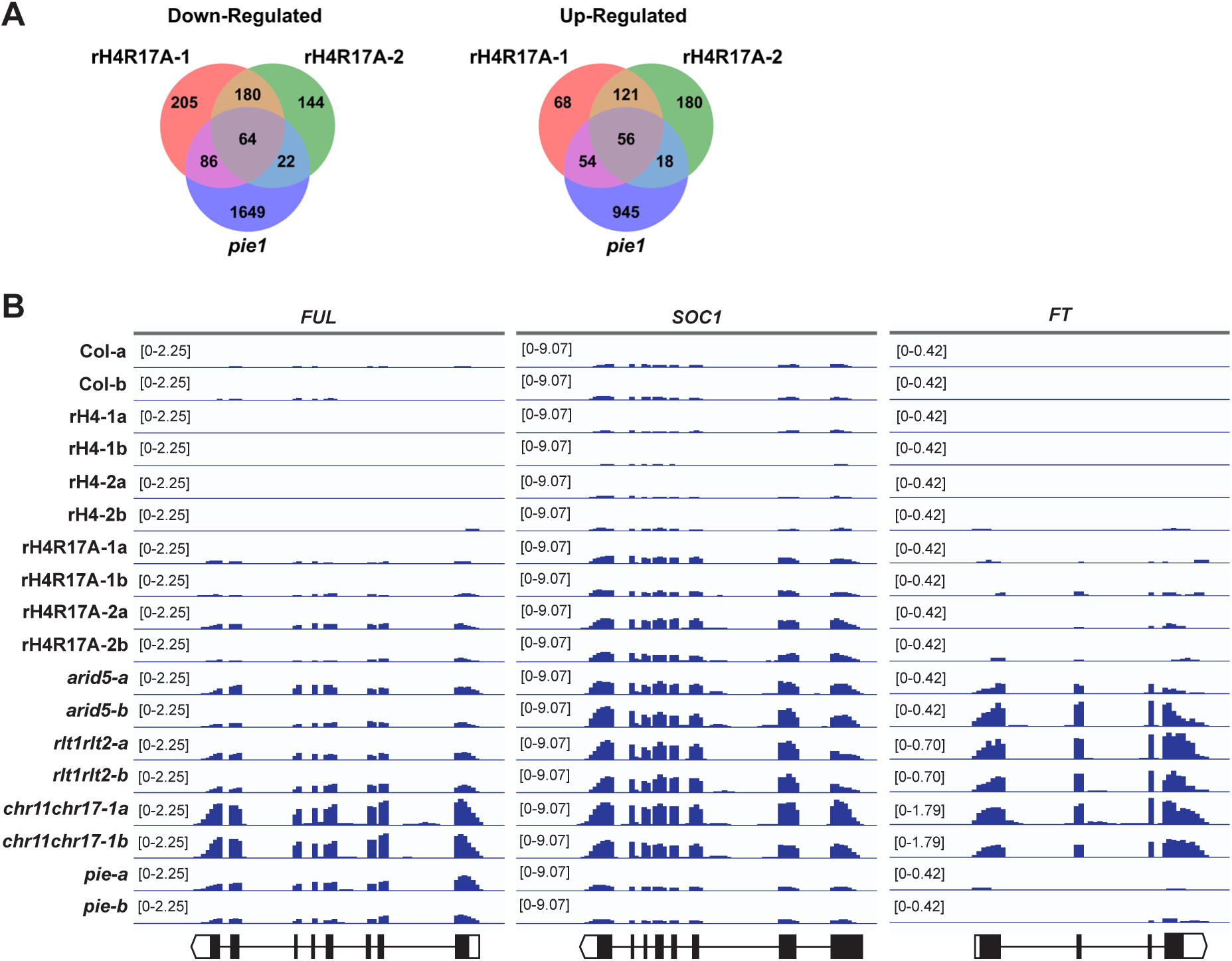
Co-regulation of gene expression observed between rH4R17A and ISWI mutants. (A) Venn diagrams showing DEGs (relative to Col) identified by RNA-seq in the rH4R17A and *pie1* mutants. (B) Genome browser view of RNA-seq signals at *FUL*, *SOC1*, and *FT* in biological replicates for Col, rH4-1, rH4-2, rH4R17A-1, rH4R17A-2, *arid5, rlt1/2, chr11/17,* and *pie1* plants. Diagrams of genes shown at the bottom, with white boxes, black boxes, and black lines representing untranslated regions, exons, and introns, respectively.

**Supplemental Figure 7:**
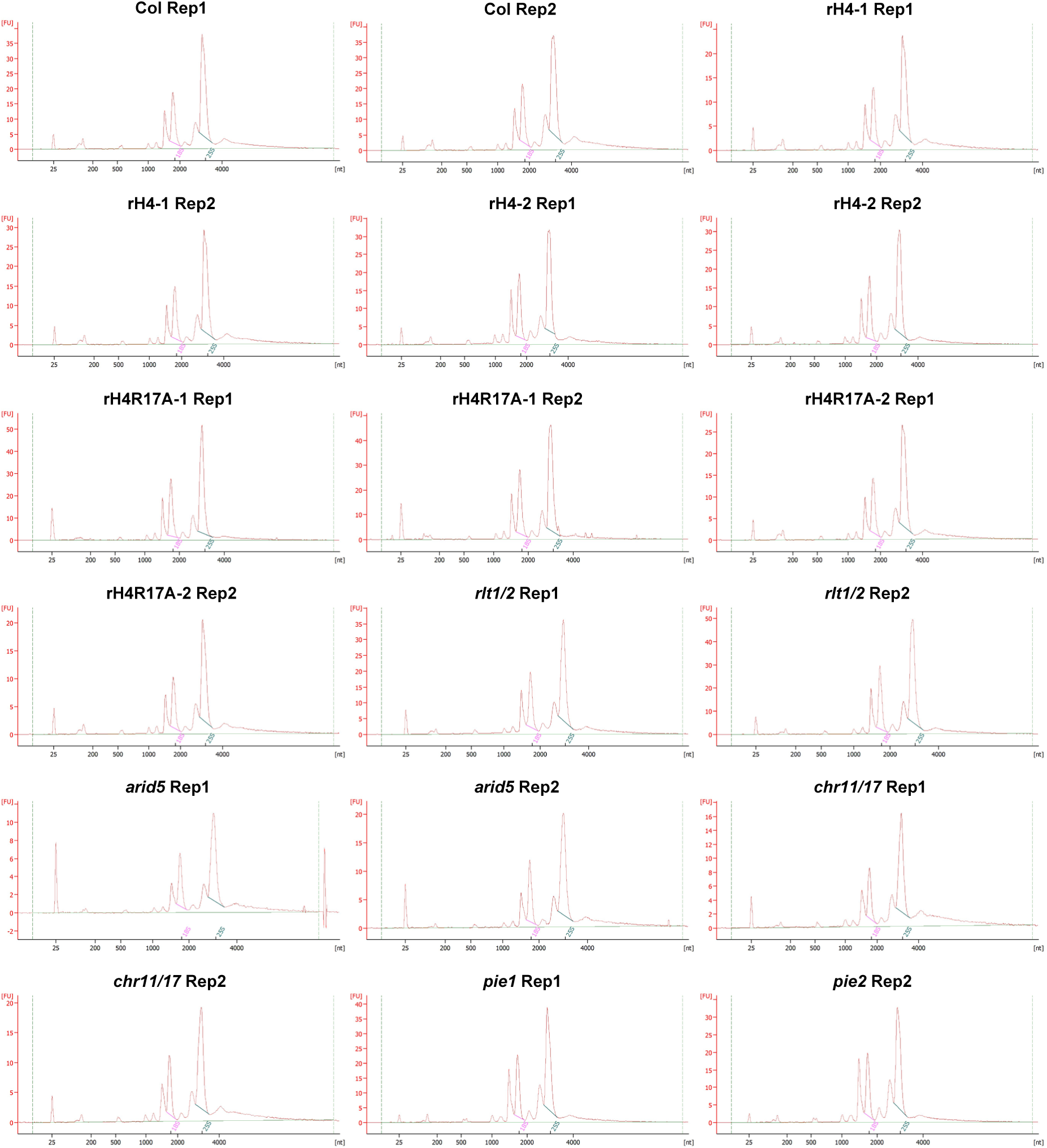
Bioanalyzer electropherograms of RNA-seq replicates. Agilent Bioanalyzer 2100 electropherograms for RNA-seq replicates of Col, rH4-1, rH4-2, rH4R17A-1, rH4R17A-2, *rlt1/2, arid5, chr11/17,* and *pie1*.

**Supplemental Figure 8:**
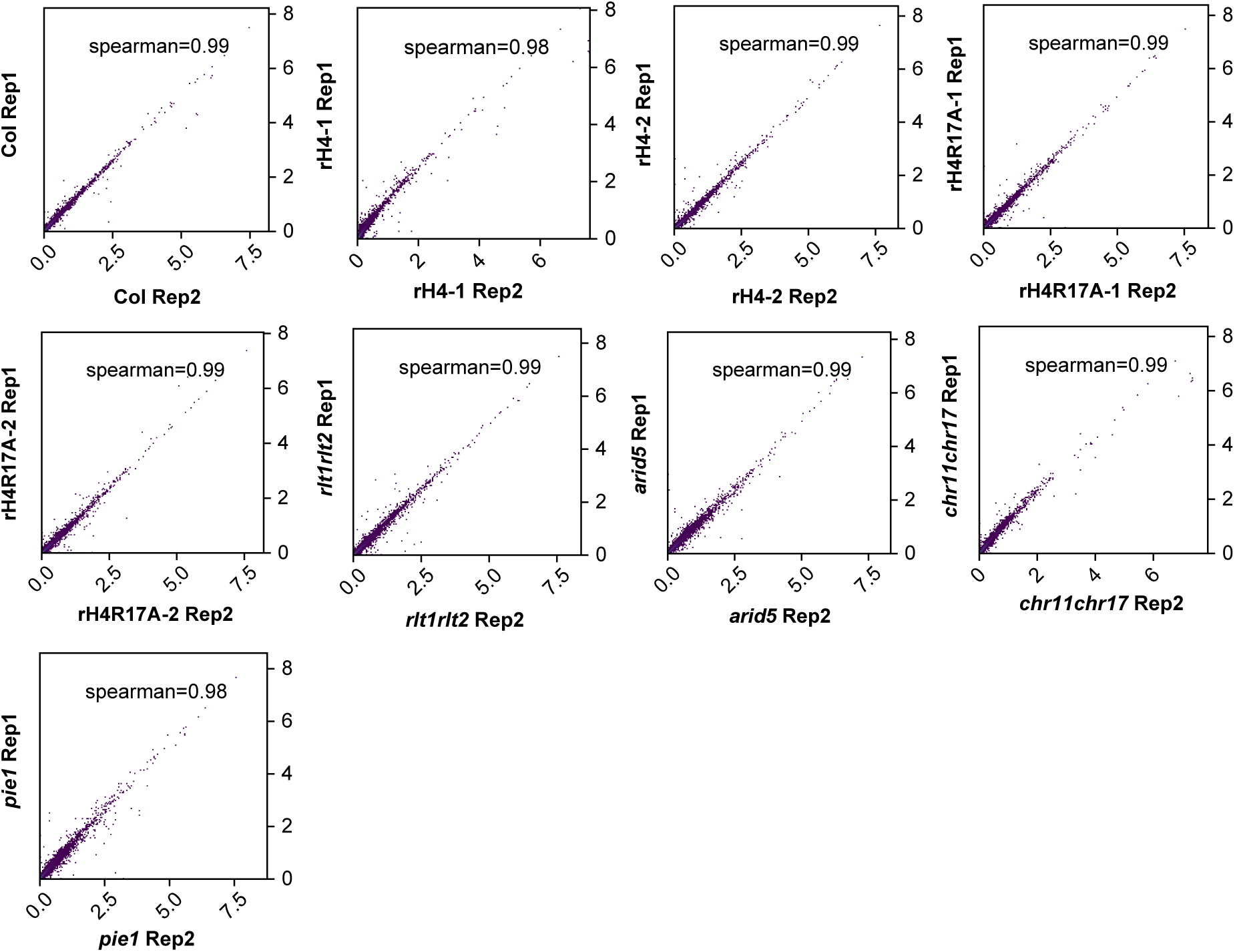
Spearman correlation of RNA-seq replicates. Spearman correlation coefficient analysis for RNA-seq replicates of Col, rH4-1, rH4-2, rH4R17A-1, rH4R17A-2, *rlt1/2, arid5, chr11/17,* and *pie1*.

**Supplemental Figure 9:**
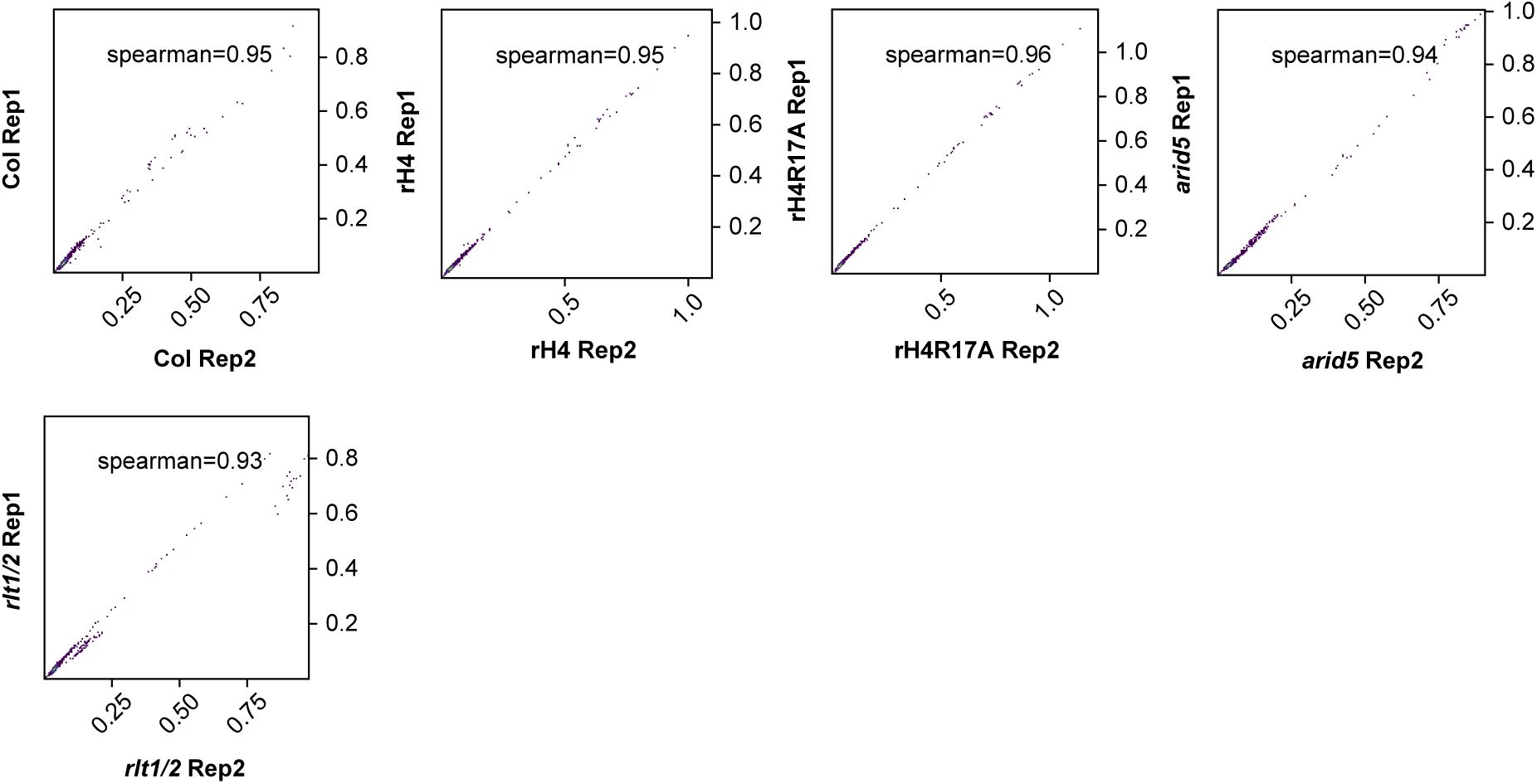
Spearman correlation of MNase-seq replicates. Spearman correlation coefficient analysis for MNase-seq replicates of Col, rH4, rH4R17A, *arid5,* and *rlt1/2*.

**Supplemental Figure 10:**
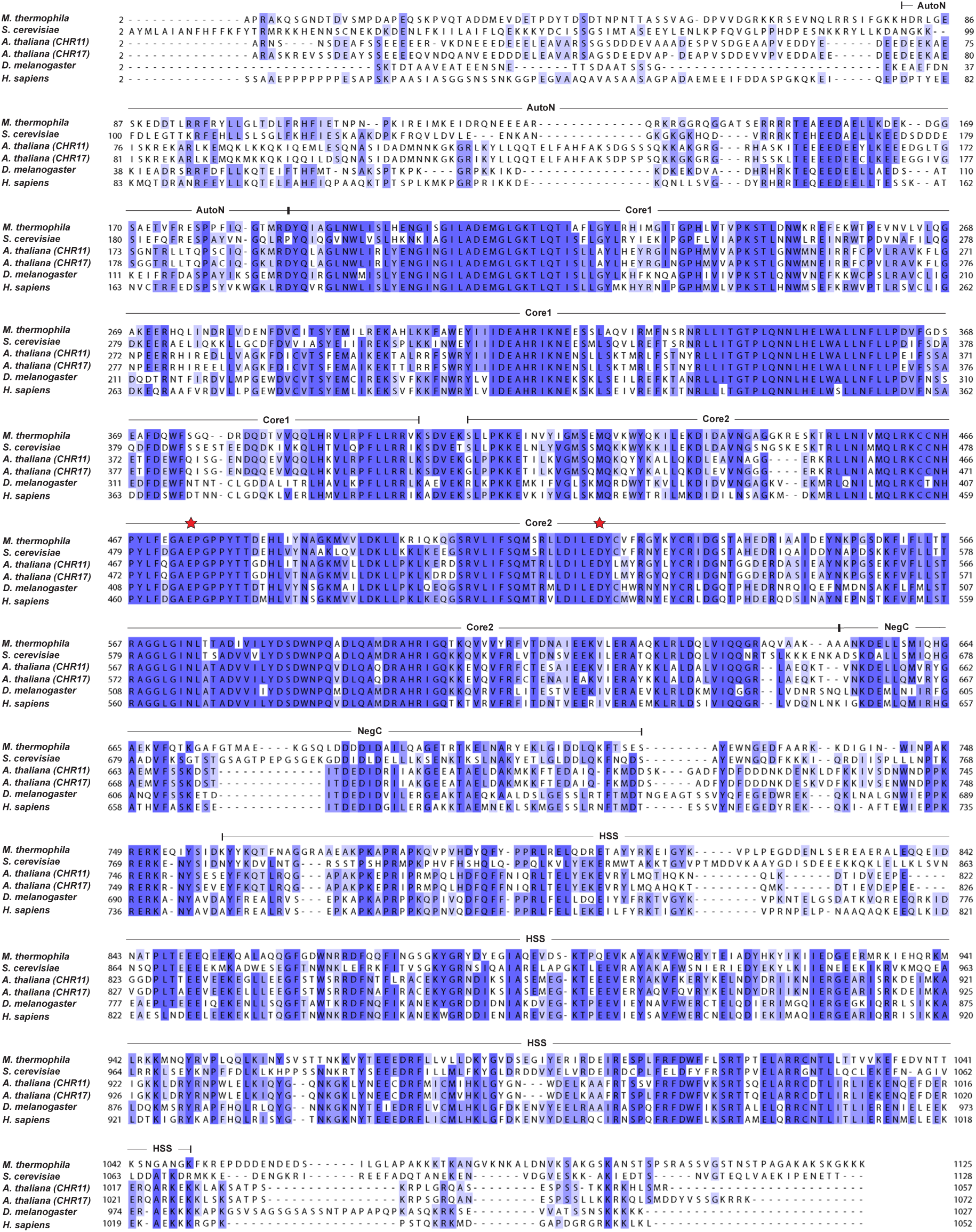
Conservation of ISWI proteins. Multiple sequence alignment of ISWI proteins performed with Clustal Omega. Protein sequences were obtained from UniProt and correspond to the following accession numbers: *Myceliophthora thermophila*: G2QFM3, *Saccharomyces cerevisiae*: P38144, *Arabidopsis thaliana* (CHR11): F4JAV9, *Arabidopsis thaliana* (CHR17): F4JY25, *Drosophila melanogaster*: Q24368, *Homo sapiens* (SNF2H): O60264. Darker shading indicates higher similarity between residues. Red stars above a.a. indicate the residues implicated in binding H4R17 on the second RecA-like ATPase core domain (core2) identified in *Myceliophthora thermophila* (Yan et al., 2016) and *Saccharomyces cerevisiae* (Yan et al., 2019). Protein domains are assigned as reported in a previous study (Yan et al., 2016). HSS: HAND–SAND–SLIDE, core1: first RecA-like domain, core2: second RecA-like domain.

**Supplemental Figure 11:**
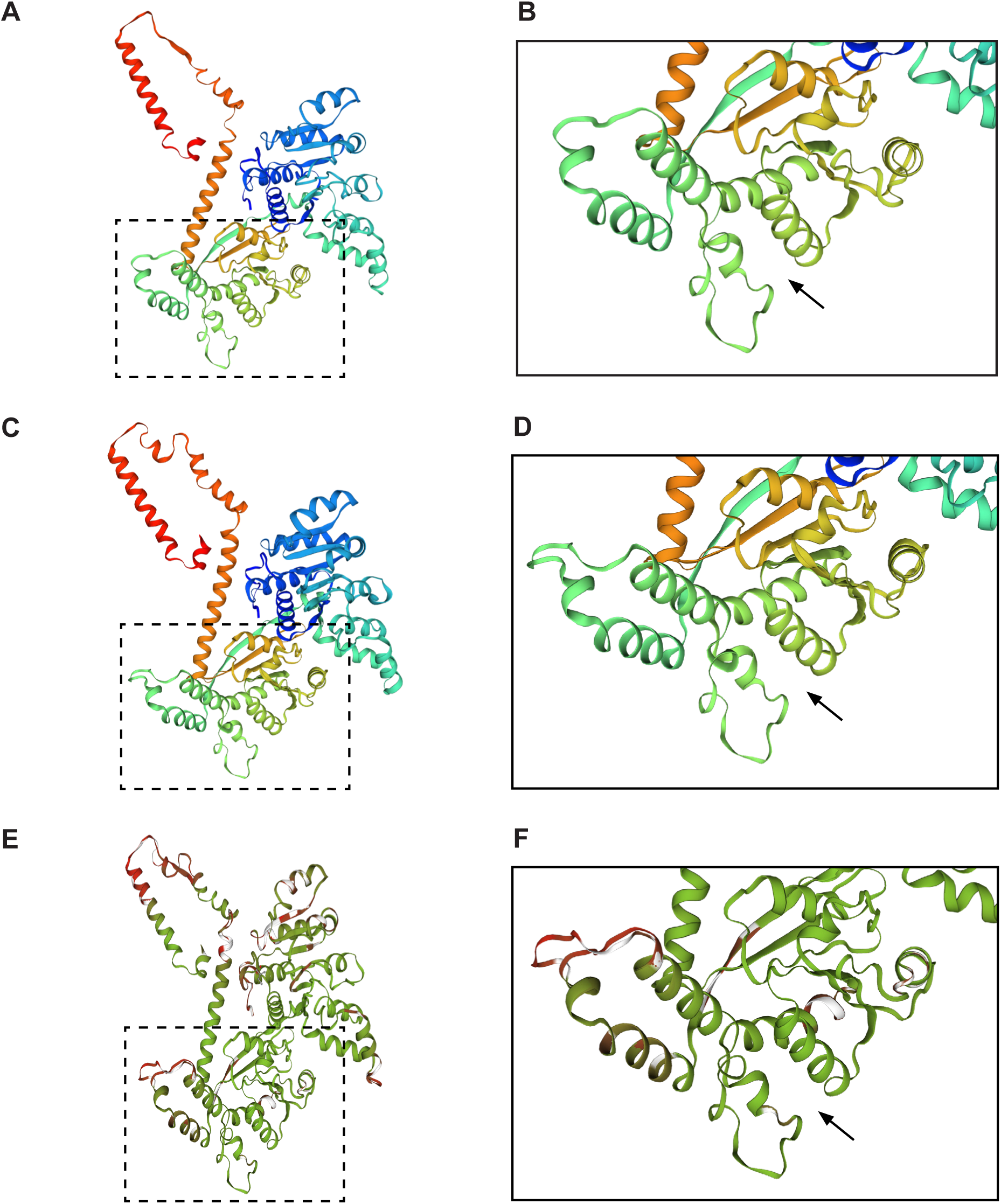
Homology model of *Arabidopsis thaliana* CHR11. (A-B) Homology model of *Arabidopsis thaliana* CHR11 a.a. 176-706. (C-D) Reference structure of *Myceliophthora thermophila* ISWI (5JXR) a.a. 173-718. (E-F) Superposition of *Arabidopsis thaliana* CHR11 and *Myceliophthora thermophila* ISWI structures with consistency color scheme (green indicates more consistent and red indicates less consistent). Black arrow denotes the predicted (*A. thaliana*) or validated (*M. thermophila*) binding pocket of histone H4 arginine 17 (Yan et al., 2016). The boxed regions are enlarged for further examinations in (B), (D), and (F).

## Notes

https://www.ncbi.nlm.nih.gov/geo/query/acc.cgi?acc=GSE190317

### Competing Interest Statement

The authors have declared no competing interest.

